# To pool or not to pool: Can we ignore cross-trial variability in FMRI?

**DOI:** 10.1101/2020.05.19.102111

**Authors:** Gang Chen, Srikanth Padmala, Yi Chen, Paul A Taylor, Robert W Cox, Luiz Pessoa

**Affiliations:** Scientific and Statistical Computing Core, NIMH, National Institutes of Health, USA; Centre for Neuroscience, Indian Institute of Science, Bangalore, India; German Center for Neurodegenerative Diseases, Magdeburg, Germany; IKND, Universität Magdeburg, Germany; Department of Psychology, Department of Electrical and Computer Engineering, Maryland Neuroimaging Center, University of Maryland, College Park, USA

## Abstract

In this work, we investigate the importance of explicitly accounting for cross-trial variability in neuroimaging data analysis. To attempt to obtain reliable estimates in a task-based experiment, each condition is usually repeated across many trials. The investigator may be interested in (a) condition-level effects, (b) trial-level effects, or (c) the association of trial-level effects with the corresponding behavior data. The typical strategy for condition-level modeling is to create one regressor per condition at the subject level with the underlying assumption that responses do not change across trials. In this methodology of *complete pooling*, all cross-trial variability is ignored and dismissed as random noise that is swept under the rug of model residuals. Unfortunately, this framework invalidates the generalizability from the confine of specific trials (e.g., particular faces) to the associated stimulus category (“face”), and may inflate the statistical evidence when the trial sample size is not large enough. Here we propose an adaptive and computationally tractable framework that meshes well with the current two-level pipeline and explicitly accounts for trial-by-trial variability. The trial-level effects are first estimated per subject through *no pooling*. To allow generalizing beyond the particular stimulus set employed, the cross-trial variability is modeled at the population level through *partial pooling* in a multilevel model, which permits accurate effect estimation and characterization. Alternatively, trial-level estimates can be used to investigate, for example, brain-behavior associations or correlations between brain regions. Furthermore, our approach allows appropriate accounting for serial correlation, handling outliers, adapting to data skew, and capturing nonlinear brain-behavior relationships. By applying a Bayesian multilevel model framework at the level of regions of interest to an experimental dataset, we show how multiple testing can be addressed and full results reported without arbitrary dichotomization. Our approach revealed important differences compared to the conventional method at the condition level, including how the latter can distort effect magnitude and precision. Notably, in some cases our approach led to increased statistical sensitivity. In summary, our proposed framework provides an effective strategy to capture trial-by-trial responses that should be of interest to a wide community of experimentalists.

## Introduction

The workhorse of functional magnetic resonance imaging (FMRI) studies is the *task design*, where it is possible to experimentally manipulate conditions to investigate the brain basis of perception, cognition, emotion, and so on. The reliability of a task-based experiment hinges on having a reasonably large number of *repetitions* associated with a condition. Such repetitions are usually termed “trials”, and each trial is considered to be an instantiation of an idealized condition. For example, in an emotion study with three conditions (positive, neutral and negative), the investigator may show 20 different human faces of each emotional valence to the subject in the scanner. From the statistical perspective, the number of trials serves as the sample size for each condition and, per the law of large numbers in probability theory, the average effect estimate for a specific condition should approximate the (idealized) expected effect with increased certainty as the number of trials grows.

Statistics lives by and flourishes in the rich variability of the data. The ultimate goal of most neuroimaging studies lies in generalizing results at the population level: the objective is to make statements that go beyond the particular samples studied. Thus, variability across samples serves as a key yardstick to gauge the evidence for the impact of experimental manipulations. More generally, in a neuroimaging study, at least four separate levels of variability are woven into the data tapestry, all of which deserve proper statistical treatment:^1^

### Cross-subject variability

Among these four levels, cross-subject variability is the easiest and most straightforward to handle. As the experimental subjects usually can be considered as independent and identically distributed, cross-subject variability is typically captured through a Gaussian distribution at the population level. In other words, each participant’s effect is considered to be drawn from a hypothetical population that follows a Gaussian distribution.

### Cross-TR variability

Because FMRI data inherently form a time series, strategies must be developed to handle the sequential dependency in the data. As the underlying mechanisms of BOLD response are not fully understood, the current models cannot exhaustively account for various effects and confounds; thus, temporal structure remains in the model residuals. The awareness of this issue has indeed generated various strategies of autoregressive (AR) modeling to tackle it.

### Cross-region variability

Multiplicity is an intrinsic issue of the massively univariate approach adopted in neuroimaging with voxels or regions treated as independent units. Various strategies have been developed, including cluster-based inferences, random field theory and permutation-based methods. At the level of regions, we recently proposed an integrative approach that handles cross-region variability with a Bayesian multilevel (BML) model that dissolves the conventional multiplicity issue (Chen et al., 2019a, 2019b).

### Cross-trial variability

Until recently, trial-by-trial response variability had received little attention (Westfall et al., 2017; Yarkoni, 2019). That is, traditional FMRI paradigms include repetitions of trials for the purpose of providing a more reliable estimate of condition-level response, and the variability is ignored. The central objective of the present study is to develop a multilevel framework to effectively handle this source of variability (along with the other sources of variability, above) in a computationally scalable approach.

What is trial-by-trial variability? Clearly, multiple sources contribute to cross-trial fluctuations, although these are rather poorly understood. When the fluctuations are of no research interest, they are often treated as *random noise* under the assumption that the “true” response to, say, a fearful face in the amygdala exists, and deviations from that response constitute random variability originating from the measurement itself or from neuronal/hemodynamic sources. Consider a segment of a simple experiment presenting five faces (Fig.1). In the standard approach, the time series is modeled with a single regressor that takes into account all face instances (Fig. 1a,b). The fit, which tries to capture the mean response, does a reasonable job at explaining signal fluctuations. However, the fit is clearly poor in several places (Fig. 1c). Traditional FMRI paradigms would ignore this variability across trials; in the present study, we propose to explicitly account for it in the modeling.

**Figure 1:**
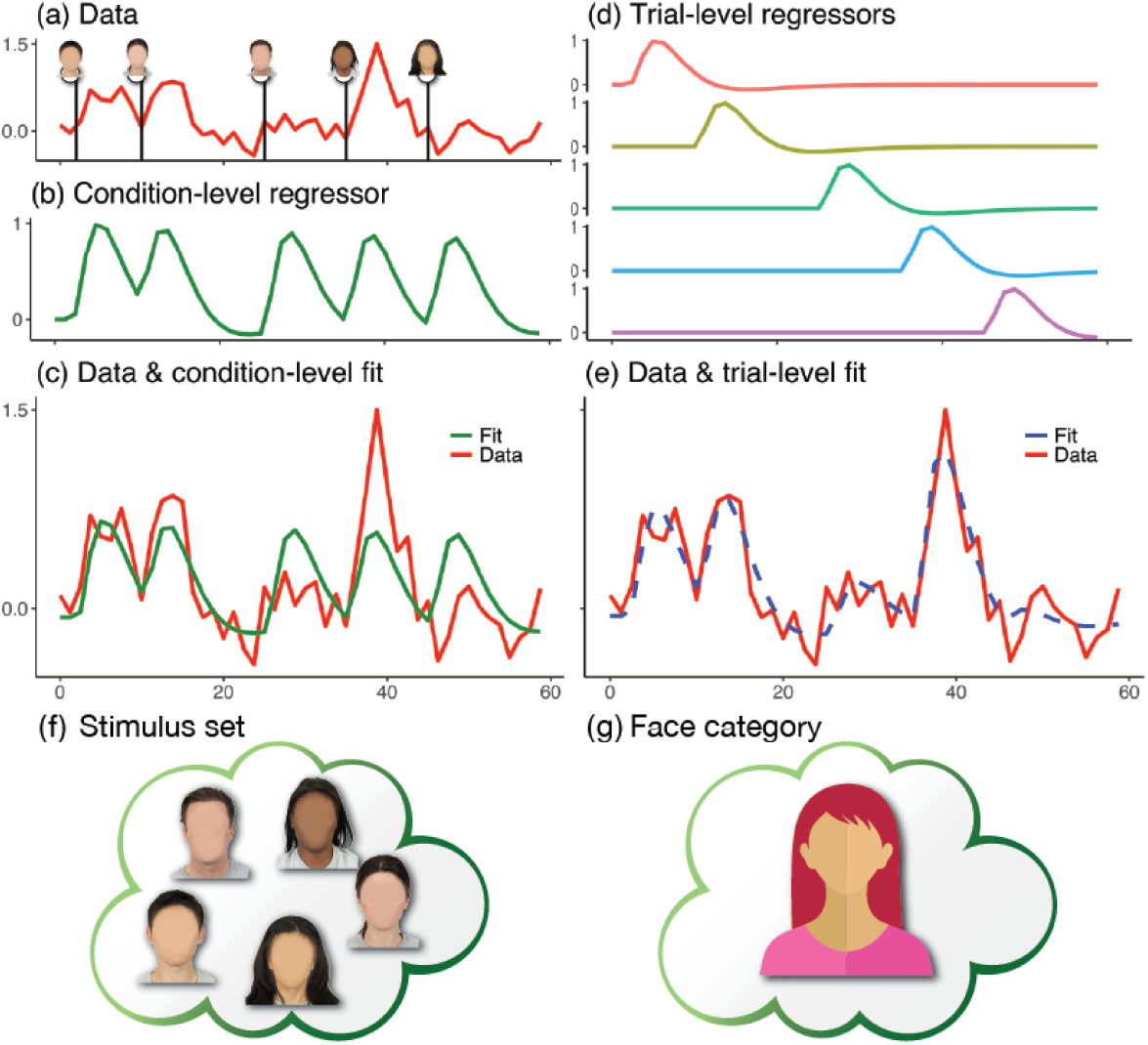
Time series modeling in neuroimaging. Consider an experiment with five face stimuli. (a) Hypothetical times series (scaled by mean value) is shown at a brain region associated with five stimuli. (b) The conventional modeling approach assumes that all stimuli produce the same response with one regressor. (c) An effect estimate (in percent signal change or scaling factor for the regressor (b)) is associated with the fit (green) at the condition level. (d) An alternative approach models each stimulus separately with one regressor per stimulus. (e) Trial-level modeling provides an improved fit (dashed blue). (f) The set of five stimuli (specific faces, blurred for privacy only) serves as a representation of and potential generalization to a condition category (face). (g) As described in the paper, trial-level estimates can be integrated via partial pooling such that inferences can be made at the general category level.

Why is it important to properly account for cross-trial variability? Under the condition-level modeling utilized in standard data analysis, trial-by-trial fluctuations are flatly swept under the rug of the model residuals, creating at least three problems:

1. an unrealistic assumption of “fixed” responses across trials;
2. the loss of hierarchical structure across the two different levels – trial and TR – of data variability;
3. the inability to legitimately generalize from the confine of specific trials (e.g., 5 neutral faces from a given stimulus dataset, Fig. 1f) to the condition category (e.g., neutral face, Fig. 1g).

This last point means that, strictly speaking, the domain of generalizability of experiment is the set of trials, which is clearly not the way experimentalists interpret their findings. If one adopts a principled trial-level modeling as developed here (see also an earlier work on trial-level modeling via a different approach, Westfall et al., 2017; we discuss the methodology differences and outcomes in our approach, below), trial-based regressors can be utilized to capture trial-by-trial fluctuations, thereby potentially capturing overall signal fluctuations better (Fig. 1d,e). Importantly, as the varying trial response is explicitly accounted for, inferences can be made at the desired level (e.g., neutral face, Fig. 1g).

There are at least three major instances where the investigator is actually interested in trial-by-trial variability. First, and perhaps most common, an investigator may associate cross-trial fluctuations to behavior. For example, trial-level effects can be associated with success/failure in task performance (Ress et al., 2000; Pessoa et al., 2002; Sapir et al., 2005; Lim et al., 2009). Second, correlation analyses can be established in a trial-by-trial fashion, at times called “beta series correlation” (Rissman et al., 2004). Third, trial-level responses are also used for prediction purposes, including multivoxel pattern analysis (MVPA), support vector machines (SVM), reinforcement learning and neural networks more generally. Our present research goal is to develop an adaptive methodology that can capture cross-trial fluctuations effectively, thus allowing them to be applied to the cases above as well as typical population-level analysis. Indeed, while we illustrate the approach with behavioral and FMRI data, it can equally be applied to MEG, EEG and calcium imaging data, and to several other paradigms—this methodology is quite general.

### Conventional time series modeling strategies

The conventional whole-brain voxel-wise analysis adopts a massively univariate approach with a two-level procedure: the first is at the subject level, and the second at the population level.^2^ The split between these two levels is usually due to two reasons. One is model complexity: because of idiosyncrasies across subjects (e.g., different trial effects and confounds, varying AR structures), it is generally unwieldy to integrate all subjects into a single model. The second consideration is practicality. In particular, it is computationally impractical to solve one “giant”, integrative model even if one could build it.

The statistical model at the subject level is time series regression^3^ solved through generalized least squares (GLS). The preprocessed EPI data *y*_*k*_ is fed into a time series regression model as the response variable on the left-hand side,

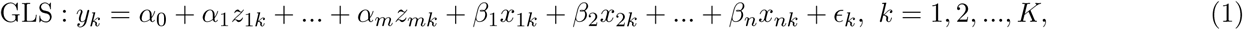

where *k* indexes discrete time points, and the residuals _*k*_ are assumed to follow a Gaussian distribution. Between the two sets of regressors, the first set *z*_*ik*_ (*i* = 1, 2, .., *m*) contains various covariates including slow drifts (e.g., polynomial or sine/cosine terms associated with low-frequency signals), head-motion variables, outlier censoring and physiological confounds such as cardiac and breathing effects, while the second set *x*_*jk*_ (*j* = 1, 2, .., *n*) is associated with the experimental conditions. Correspondingly, there are two groups of effect parameters: the first set *α*_*i*_ (*i* = 1, 2, .., *m*) is usually of no interest to the investigator while the second set *β*_*j*_, *j* = 1, 2, …, *n* is the focus of specific research questions.

The construction of condition regressors *x*_*jk*_ (*j* = 1, 2, .., *n*) in the GLS model (1) largely depends on the research focus. For most investigations, the interest is placed on the effects at the condition level, and the trials of each condition are treated as multiple instantiations of the event of interest. While various approaches are adopted to construct the condition regressors *x*_*jk*_, they are typically treated with the assumption that the response magnitude remains the same across all trials of a condition (Fig. 1b,c). Specifically, one regressor per condition is constructed through the convolution of the individual trial duration with a fixed-shape hemodynamic response function.^4^ Note that the fixed-response-magnitude approach can be relaxed in certain scenarios. For example, one may modulate the trial-level response by creating another regressor through auxiliary information (e.g., reaction time (RT)). At present, we will focus on an alternative approach: to capture the trial-level effects, one feeds one regressor per trial to the GLS model (1); trial-level modulation, if desired, will be performed at the population level.

Another complexity involves the residuals ϵ_*k*_ in the GLS model (1). If the residuals are white (i.e., no autocorrelation), time series regression can be numerically solved through ordinary least squares (OLS) or maximum likelihood. However, it has been long recognized that temporal correlation structure exists in the residuals (e.g., Bullmore et al., 1996) because some components in the data either are unknown or cannot be properly accounted for. Failure to model the autocorrelation may lead to inflated reliability (or underestimated uncertainty) of the effect estimates. Three strategies that utilize GLS have been proposed to improve the model by characterizing the temporal correlations in the residuals ϵ_*k*_. First, an early approach was to characterize the autocorrelation with a uniform first-order AR model for the whole brain (Friston et al., 2002). Second, a localized AR(1) model was developed later so as to consider neighboring voxels within each tissue type (Woolrich et al., 2001). Third, an even more flexible approach was created using an autoregressive moving average ARMA(1,1) structure that accommodates the model at the voxel level through the program 3dREMLfit in AFNI (Chen et al., 2012). A recent comparison study has shown that the performances of the three methods match their respective modeling flexibility, complexity and adaptivity (Olszowy et al., 2019).

The conventional modeling of condition regressors in the GLS model (1) can be further extended. For one, we can take inspiration from typical population-level analysis, which includes a term for each subject so that subject-specific effects are properly accounted for. The same approach can be adopted at the trial level to account for trial-by-trial variability. In particular, the assumption that all the trials of a given condition share the same brain response magnitude should be viewed skeptically (Fig. 1d,e). Critically, from a modeling perspective, treating trials as “fixed effects” is tantamount to limiting the focus of the study to the trial instantiations employed, potentially exaggerating the statistical evidence and foiling the validity of the experimenter’s goal to generalize from the particular samples used (e.g., specific faces utilized in the experiment) to the generic level (e.g., human faces in general). Needless to say, the latter generalization is taken for granted in neuroimaging studies. Here, we argue that the modeling strategy adopted should address this issue head on, and we demonstrate a trial-level modeling approach to achieving this goal.

## Methods

### Perspectives on trial-level modeling

Our motivation is to directly model trial-level effects at the subject level and to account for across-trial variability at the population level. In doing so, the conventional assumption of constant response across trials is abandoned in light of the following two perspectives.

#### 1) Research focus

Depending on the specific research hypothesis, one may be interested in: (a) trial-level effect estimates for each subject, so that those effects can be utilized for predictions or correlativity among regions; (b) association of trial-level effects with behavioral data; or (c) condition-level effects. We will focus on the latter two which involve population-level analysis.

#### 2) Modeling perspective

The BOLD response magnitude varies across trials, but what is the nature of the trial-to-trial fluctuations? There are three modeling strategies depending on the ultimate research goal, mapping to three different data pooling methods (Chen et al., 2019b). The first, commonly adopted approach assumes that the underlying BOLD response does not change from trial to trial and that the observed fluctuations are noise or random sampling variability (Fig. 1b,c). Thus, the average response is estimated across trials to represent condition-level effects. This approach can be considered to be *complete pooling*, where all the “individuality” of trials is ignored in the model. Technically, the approach precludes generalization to the trial category in question, and does not allow extending one’s conclusions from the specific trials used in the experiment to situations beyond the trials employed (Yarkoni, 2019). In contrast, it is possible to adopt a *no pooling* strategy at the subject level, and estimate each trial’s response separately (Fig. 1d,e); in other words, each trial is fully unique and assumed to be unrelated to other trials. Between the two extremes, a middle ground can be taken at the population level such that the cross-trial variations are considered as random samples of the condition-level effect (cf. subjects as samples of an idealized population). This characterization of randomness allows the investigator to make the generalization from the specific trials instantiated in the experiment to the *concept* of a condition category, the idealized population from which trials are envisioned to be random samples. With such *partial pooling* approach, information can be loosely but meaningfully shared across trials.

Neuroimaging is no stranger to dealing with the three pooling methods. In fact, the issue about cross-trial variability basically runs parallel to its cross-subject counterpart. The typical split between the subject- and population-level analyses means that a no-pooling strategy is adopted at the individual subject level in the sense that each subject is assumed to have unique response effects; then partial-pooling is typically followed up at the population level with a Gaussian distribution for cross-subject variability. In the early days, there were even choices between fixed-versus random-effects analysis at the population level; such a comparison is just another way to elaborate the differences between complete and partial pooling. Today, complete pooling for cross-subject variability (or fixed-effects analysis) is typically considered unacceptable (leading to paper rejection!), and the adoption of partial pooling (or random-effects analysis) at the population level is routine practice. It is exactly the same underlying rationale that we wish to address in the context of cross-trial variability, thus we believe there are no legitimate reasons preventing the analyst from the adoption of a more general pooling methodology.

### Population analysis through trial-level modeling

We start with a linear mixed-effects (LME) platform for population analysis. The model incorporates trial-level effect estimates *y*_*st*_ under one condition from individual subjects based on the GLS model (1),

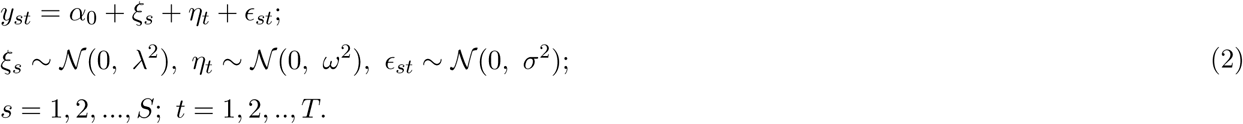

The indices *s* and *t* code subjects and trials, respectively; *α*_0_ is the intercept that embodies the overall effect at the population level; *ξ*_*s*_ and *η*_*t*_ represent cross-subject and cross-trial effects (random effects); ϵ_*st*_ is the residual term and usually assumed to follow a Gaussian distribution. When explanatory variables are involved (e.g., between- and/or within-subject variables), the model can be naturally extended by augmenting the intercept term *α*_0_. The LME framework (2) with a crossed or factorial random-effects structure can be numerically analyzed by, for example, the program 3dLMEr^5^ in AFNI (Cox, 1996) at the whole-brain voxel-wise level.

How does the conventional approach compare to the LME formulation for trial-level modeling? The former can be conceptualized as a dimensional reduction step of so-called “summary statistics”, a common practice in neuroimaging. With the assumption that the effects from the *T* trials follow Gaussian distribution, the conventional approach essentially reduces the whole distribution, one-dimensional curve, with one number, its mean. Such dimensional reduction may substantially decreases the amount of data and simplifies the model at the population level, information loss or distortion naturally becomes a legitimate concern.

What is the exact direct impact when cross-trial variability is ignored? In the conventional approach trial effects are obviously not modeled at the subject level. To a first approximation, the condition-level effect can be conceptualized as the arithmetic mean of the trial-level effect estimates 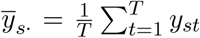. When no trial-level information is available, the LME model (2) simply reduces to the conventional Student’s *t*-test for the condition-level effects 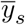:

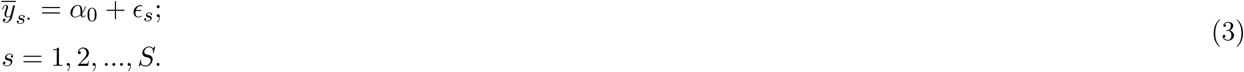

With assumption of an identical and independent distribution of cross-trial effects 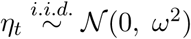, the missing component of the cross-trial effects *η*_*t*_ in the GLM (3) relative to the LME counterpart (2) means that the variability,

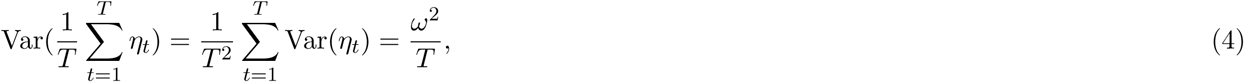

is not accounted for in the condition-level approach.^6^ Notice that this variability depends on and is sensitive to *T*, the number of trials. Therefore, if *T* is large enough, the variability may become inconsequential, in which case complete pooling could be justified as a reasonable approximation. However, given that the number of trial repetitions is relatively small, such scenario appears unrealistic. Accurately mapping the data hierarchy and explicitly characterizing cross-trial variability, as represented by the trial-specific terms *η*_*t*_ and its distribution *𝒩* (0, *ω*^2^) in the LME model (2), legitimizes the generalizability from the specific trials to a general category.

The early history of FMRI data analysis sheds some light on the issue of ignoring cross-trial variability. In the early 2000s, there was an active debate in the field about population-level analysis, specifically between aggregating cross-subject estimates through complete pooling (or fixed-effects analysis) versus partial pooling (or random-effects analysis). Presently, it is clear that ignoring cross-subject variability leads to underestimated uncertainty and inflated statistical evidence (Penny and Holmes, 2007), and there is now consensus that partial pooling/random-effects is necessary to draw adequate inferences at the population level. Our investigation of cross-trial variability can be conceptualized along the same lines as the older debate, but now considering another source of variability, namely, trials. Although the analytical aspects are now more complex given the additional dimension of effect decomposition (i.e., trials), we believe the consequences (e.g., effect inflation) of ignoring cross-trial variability are similar to those of ignoring cross-subject variability (e.g., Baayen et al., 2008; Westfall et al., 2017).

We reiterate that it is through the explicit capture of cross-trial variability that provides a solid foundation for generalization. As a routine practice, nowadays cross-subject variability is properly accounted for at the population level, and such accountability is evidenced in conventional models as simple as Student’s *t*-tests, GLM and AN(C)OVA, or as the subject-specific terms *ξ*_*s*_ and their distribution *𝒩* (0, *λ*^2^) in the above LME platform (2). However, the same rationale has not been adopted and applied to the cross-trial variability, even though the adoption of many exemplars of a condition in experimental designs is intended for generalization.

One potential improvement of the LME model (2) is the incorporation of effect precision. The subject-level effect estimates (e.g., *y*_*st*_ in the LME model (2)) from the GLS model (1) are estimated, naturally, with some degree of uncertainty (embodied by the standard error, 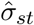). As the whole analysis pipeline is broken into the two levels of subject and population, theoretically it is desirable to explicitly incorporate the reliability information of the effect estimates into the population-level model so that the information hierarchy would be largely maintained. In standard practices, subject-level standard error is usually ignored at the population level; such practice assumes that the uncertainty is either exactly the same across subjects or negligible relative to the cross-subject variability (Chen et al., 2012). To address this shortcoming, some population-level methods have been developed to incorporate both effect estimates from the subject level and their standard errors (Worsley et al., 2002; Woolrich et al., 2004; Chen et al., 2012). However, this integration approach has not gained much traction in practice due to its small potential gain (Mumford et al., 2009; Chen et al., 2012; Olszowy et al., 2019). Within the LME framework, unfortunately there is no easy solution to consider these standard errors. In contrast, taking them into account is a natural component of BML modeling, and we will explore the role of precision information in the current context of trial-level modeling.

Another possible improvement of the LME model (2) is outlier handling. Due to the substantial expansion in the number of regressors involved in trial-level modeling, the chance of having outlying effect estimates cannot be ignored. However, it is a challenge to handle outliers and data skew within the LME framework. A typical approach is to set hard bounds, thus constraining data to a predetermined interval in order to exclude outliers. In contrast, by adopting a BML framework, outliers can be accommodated in a principled manner with the utilization of non-Gaussian distributions for data variability.

### Handling behavioral covariates and nonlinearity

Trial-level modeling can be extended to incorporate behavioral variables. In conventional approaches, the association between trial-level effects and behavior can be modeled by creating a modulatory variable at the subject level. Accordingly, instead of one, two regressors are constructed per condition. The first is the typical regressor for the average condition effect (here, the behavioral measure is considered at a center value, such as the subject’s mean). The second regressor codes, for example, for the linear relationship between BOLD response and the behavioral measure. When trial-level effects are directly estimated at the subject level, the following LME can be adopted at the population level:

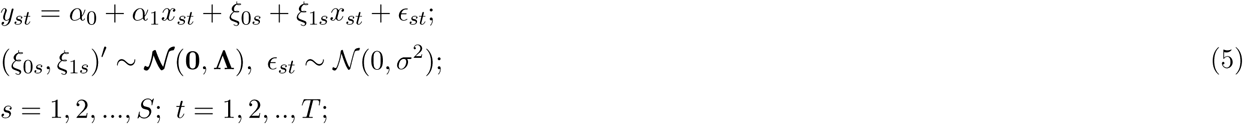

where *x*_*st*_ is the behavioral measure of the *s*th subject at the *t*th trial. The parameters *α*_0_ and *ξ*_0*s*_ are population- and subject-level intercepts, respectively. The effect of the behavioral variable *x*_*st*_ on the response variable *y*_*st*_ is captured through the slope parameter *α*_1_ at the population level, while its subject-level counterpart is characterized by the slope parameters *ξ*_1*s*_. The 2 × 2 variance-covariance matrix **Λ** reflects the relationship between the subject-level intercept *ξ*_0*s*_ and slope *ξ*_1*s*_.

The modeling of behavioral covariates can be altered to relax the linearity assumption. Polynomials (e.g., quadratic terms) can be used but still require some extent of prior knowledge and assumption about the relationship. Alternatively, we can adopt smoothing splines with a set of basis functions defined by a modest sized set of knots. For example, we can use penalized cubic smoothing splines *s*(*·*) to achieve a counterbalance between the goodness of fit and the curvature or wiggliness measured by the integrated square of second derivative (Wood, 2017):

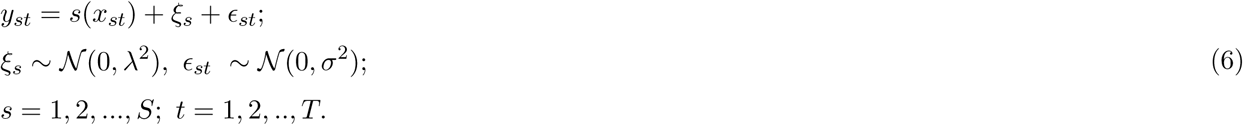

The approach of smoothing splines can be conceptualized as an adaptive and calibrating process. On one hand, we could simply adopt a naive fit with a straight line to the data; on the other hand, we could fully rely on the data and trace faithfully each data point, regardless of the roughness of the fitting curve. Between these two extremes, we intend to learn from the data by searching for a middle ground through an adaptive process of partial pooling with the imposition of curve smoothness.

### Alternative trial-level modeling

Previously, Westfall et al. (2017) proposed an integrative approach to address the trial-level generalization problem. They relaxed the assumption of fixed BOLD response across trials and directly modeled trial-to-trial fluctuations with the presumption of an AR structure in the data *y*_*sk*_ as opposed to the residuals in addition to a few oversimplifications. Furthermore, both the subject and population levels were merged into one model. In the following description, we have slightly modified and generalized their original notation with data of *I* experimental conditions from *S* subjects,

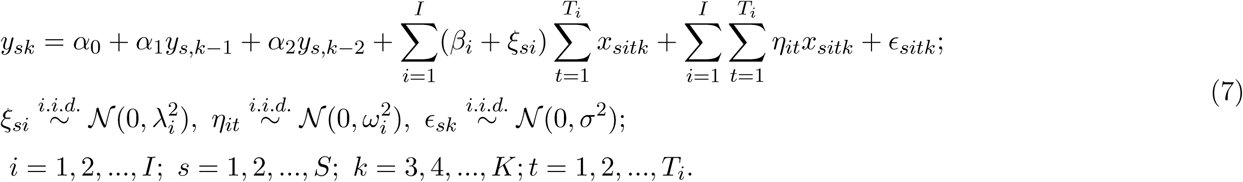

Indices *s, i, t* and *k* code subject, condition, trial and time, respectively; *T*_*i*_ is the number of trials for the *i*th condition; *α*_0_ is the overall intercept; 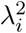 and 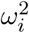 characterize the cross-subject and cross-trial variability, respectively, for the *i*th condition; *x*_*sitk*_ is the trial-level regressor; the effect associated with the regressor *x*_*sitk*_ of the *i*th condition is partitioned into two components, *β*_*i*_ for the average (fixed) component across all trials and *ξ*_*si*_ for the subject-specific (random) component; *η*_*it*_ represents the cross-trial (random) effect shared by all subjects; _*sitk*_ is the residual term with the assumption of white noise (no serial correlation) and variance *σ*^2^. Note that an AR(2) structure with two parameters *α*_1_ and *α*_2_ is explicitly modeled with lagged effects as regressors, instead of being embedded in the residuals as is typically practiced in the field (Woolrich et al., 2001; Worsley et al., 2002; Chen et al., 2012). All random effects and residuals are assumed Gaussian. In addition, likely for computational simplifications, the intercept *α*_0_, AR effects *α*_1_ and *α*_2_, cross-trial effect *η*_*it*_ are assumed to be the same across subjects. Due to the unavailability of numerical implementations and the intractable computational cost, the above LME model was solved at the region level in Westfall et al. (2017) through the NiPyMC Python package. Finally, they focused solely on conventional statistical evidence and its dichotomization (i.e., thresholding), whereas we wish to consider both effect magnitude and the associated statistical evidence through a more continuous view of statistical support (Chen et al., 2017).

### Trial-level modeling and study goals

We will use an FMRI dataset to demonstrate our trial-level modeling framework that blends in well with the current analytical pipeline. At the subject level, the effect estimate at each trial is obtained with no pooling through the GLS model (1), with the temporal correlation in the residuals captured via an ARMA(1,1) structure. At the population level, in parallel to cross-subject variability, the trial-level effects are modeled through partial pooling to address the following question: What are the differences and consequences compared to the conventional approach of complete pooling? The common practice in neuroimaging is largely limited on statistical evidence followed by artificial dichotomization; thus, relatively little attention is paid to effect magnitude. For example, Westfall et al. (2017) reported substantially inflated statistical values (1.5-3.0 times) when complete pooling was adopted. Here, we wish to explore whether we could develop a computationally economical approach to incorporating trial-level effects while emphasizing the impact of trial-level modeling on both effect estimate and its uncertainty. Overall, the issues that we want to raise and explore include:

1. extent of cross-trial variability;
2. variability of autocorrelation structure in the subject-level residuals;
3. impact of directly modeling the autocorrelation with lagged effects;
4. cross-trial fluctuations as an indication of synchrony among regions;
5. importance of incorporating precision information in model formulation;
6. handling of data skew and outliers;
7. reporting full results in a comprehensive fashion.

Most of the models in this paper are under the Bayesian framework using Stan (Carpenter et al., 2017) through the R package brms (Bürkner, 2018). The choice of Bayesian modeling was made for multiple reasons, most notably its ability to incorporate multiplicity and to provide a straightforward interpretation of effect estimates through posterior distributions, instead of using point estimates and significance testing thresholding. Each Bayesian model here is specified with a likelihood function, followed by priors for lower-level effects (e.g., trial, region, subject). The hyperpriors employed for model parameters (e.g., population-level effects, variances in prior distributions) are discussed in Appendix B. Note, however, that if the ROI-related components in our models are excluded, the models can be applied at the whole-brain voxel level under the conventional LME framework.

### Trial-level modeling of FMRI data

#### Experimental data

We adopted a dataset from a previous experiment (Padmala et al., 2017). A cohort of 57 subjects was investigated in a 3T scanner. Each subject performed 4 task types,^7^ each of which was repeated across 48 trials. Each task started with a 1 s cue phase indicating the prospect of either reward (`Rew`) or no-reward (`NoRew`) for performing the subsequent task correctly. The cue was followed by a 2-6 s variable delay period. The task stimulus itself was displayed for 0.2 s. Participants had to perform a challenging perceptual task when confronted with either a negative (`Neg`) or neutral (`Neu`) distractor. The subject was then expected to respond within 1.5 s. The total 4 × 48 = 192 trials were randomly arranged and evenly divided across 6 runs with 32 trials in each run. The TR was 2.5 s. In the analyses that follow, only correct trials were employed.

We sought to investigate the interaction between motivation (reward) and emotion (distraction) in both behavior and brain data. The experiment manipulated two factors: one was the prospect (Pro) of being either rewarded or not while the other was the distractor (Dis) displayed, which was either negative or neutral. In terms of brain responses, there were six effects of interest: two cue types (`Rew`, `NoRew`) and four prospect-by-distractor task types (`NoRew_Neg`, `NoRew_Neu`, `Rew_Neg`, `Rew_Neu`) following a 2 × 2 factorial structure. In terms of behavior, the focus was the recorded trial-level RT on the same four task types. In addition, the relationship between brain response and behavioral RT was of interest. Here, variable names with a first capital letter (e.g., Pro and Dis for the manipulation factors) symbolize population effects (or fixed effects under the conventional framework), whereas those with a first lowercase letter (e.g. subj, trial and roi for subject, trial and ROI) indicate lower-level (or random) effects.

#### Behavioral data analysis

We first analyzed behavioral performance at the condition level. The success rate, rate_*ijs*_, measures the proportion of correct responses (out of 48 trials) of *s*th subject under the task of *i*th prospect and *j*th distractor. The data could be analyzed with a binomial distribution through a logistic model. However, to aid interpretability, as the number of trials of each task was reasonably large, the binomial distribution was approximated as Gaussian; thus, we opted to model the success rate data in a manner that follows the model:

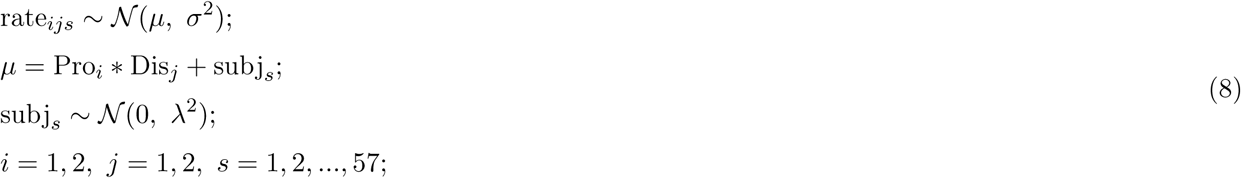

where subj_*s*_ is the subject-specific effect, Pro_*i*_ is the effect associated with the *i*th level of prospect, and Dis_*j*_ is the effect associated with the *j*th level of distractor; the expression Pro_*i*_*Dis_*j*_ is *α*_0_+Pro_*i*_+Dis_*j*_ +Pro_*i*_ : Dis_*j*_ with Pro_*i*_ : Dis_*j*_ being the second-order interaction between the two variables (borrowing the notation convention from the statistical programming language *R*).^8^ We also fitted the success measure data using a *t*-distribution instead of Gaussian in the model (8); however, the modification did not improve model fit considerably.

The response accuracy data rate_*ijs*_ is consistent with the following conclusions. Accuracy varied to some extent across the four conditions (Fig. 2a). For the main effects of the two factors (prospect and distractor, bottom two rows, Fig. 2b), the subjects had a lower response accuracy for the `NoRew` condition than `Rew`, while the accuracy for the two distractors types `Neg` and `Neu` was comparable. The overall interaction between prospect and distractor, (`NoRew`−`Rew`):(`Neg`−`Neu`), was fairly robust (top row, Fig. 2b). Specifically, the prospect effect (`NoRew`−`Rew`) was larger under the `Neg` distractor than `Neu` (second and third row, Fig. 2b) while the distractor effect (`Neg`−`Neu`) was largely in the opposite direction between the two prospects of `NoRew` and `Rew` (fourth and fifth row, Fig. 2b).

**Figure 2:**
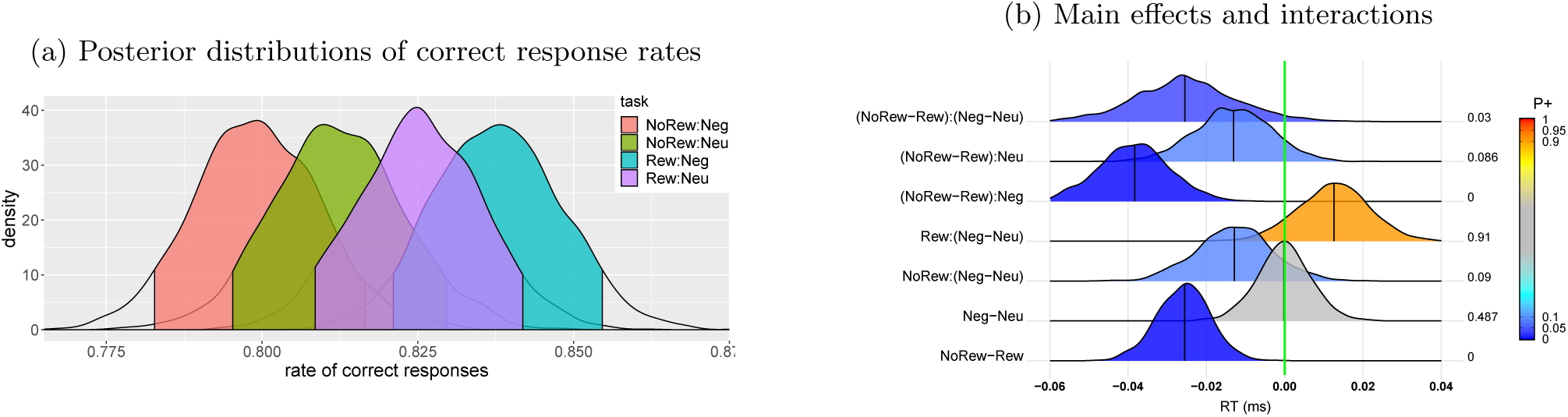
Summary of accuracy based on the BML model (8). (a) Response accuracy and the associated 95% quantile interval were estimated for each of the 4 tasks. (b) Among the posterior distributions of accuracy, the bottom two rows are the main effects while the top five rows show the interactions. At the right side of each distribution lists the posterior probability of each effect being positive, P+ (area under the curve to the right of the green line indicating zero effect), also color-coded in the distribution shading. The vertical black line under each distribution is the median (or 50% quantile). Each distribution is a kernel density estimate which smooths the posterior samples. This figure corresponds to Fig. 3B in Padmala et al. (2017).

Now we focus on the RT data at the trial level. The RT analyses had to deal with the issue of trial-versus condition-level dichotomy, illustrating the differentiation between complete and partial pooling. Across participants, the number of correct responses ranged from 28 to 47 out of 48. We constructed the following model that directly accounts for cross-trial variability,

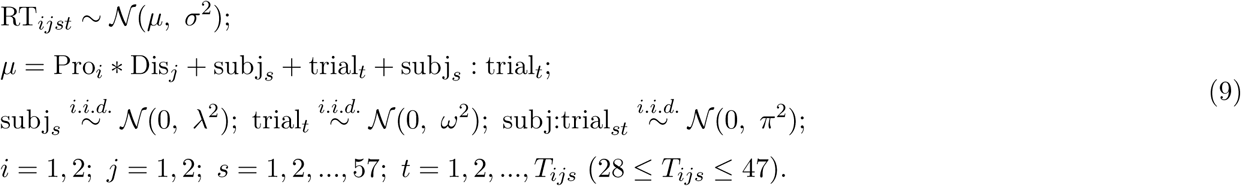

The terms Pro_*i*_ and Dis_*j*_ are the effects associated with the prospect and distractor level, respectively; subj_*s*_, trial_*t*_, and subj:trial_*st*_ are the varying effects associated with the *s*th subject, *t*th trial and their interaction, respectively; *T*_*ijs*_ is the number of correct responses of the *s*th subject during the task of *i*th prospect and *j*th distractor. Examination of the RT data indicated that the overall distribution was skewed to some extent (Fig. 3a). Thus, we explored two modified models using either a Student *t*-student or an exponentially modified Gaussian distribution (Palmer et al., 2011) to handle the skew, simply by replacing *𝒩* (*µ, σ*^2^) in the model (9) with the Student’s *t*-distribution *𝒯* (*ν, µ, σ*^2^) or *ε ℳ 𝒢* (*µ, σ*^2^, *β*), respectively, where *ν* is the parameter that codes the degrees of freedom for the *t*-distribution, and *β* is an exponential decay parameter for the exGaussian distribution. These two models produced similar effect estimates and statistical evidence. However, they provided improved fitting: the estimated degrees of freedom, *ν*, had a mean of 3.4 with 95% quantile interval of [3.1, 3.7], consistent with the skewness of the data; skewness was also accommodated by the exponential decay parameter estimate *β* = 106.19 *±* 1.81 ms.

**Figure 3:**
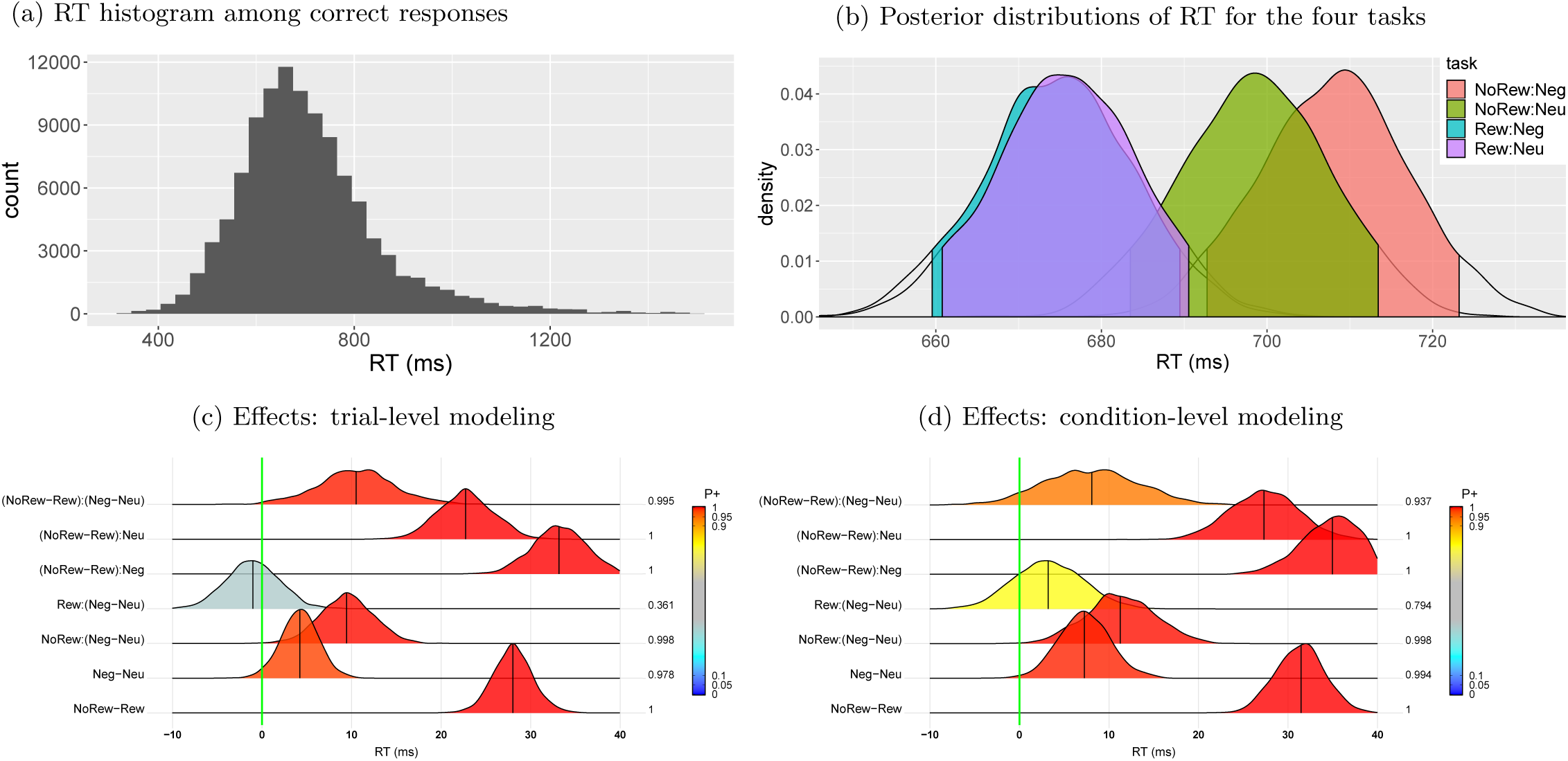
Summary of RT data based on the BML model (9) with *t*-distribution. (a) The histogram of RT among correct response trials shows the aggregated information across the trials (within [28, 47]), 4 tasks and 57 subjects (bin width: 30 ms). (b) RT and the associated 95% quantile intervals were shown for each of the 4 tasks with an overall mean of 689.3ms and s.d. of 8.8ms. (c) Among the posterior distributions based on the model (9), the bottom two rows are the main effects while the top five rows show the the interactions. At the right side of each distribution lists the posterior probability of each effect being positive, P+ (area under the curve to the right of the green line indicating zero effect), also color-coded in the distribution shading. The black vertical segment under each distribution shows the median. (d) The counterpart result of (c) based on the condition-level RT effects aggregated cross trials (corresponding to Fig. 3A in Padmala et al. (2017)).

The RT data supports the following conclusions. The posterior distribution was different among the four tasks (Fig. 3b). For the main effects of the two factors (prospect and distractor, bottom two rows, Fig. 3c), RTs were substantially shorter during `Rew` trials, and `Neg` distractors robustly slowed down behavior. The overall interaction between prospect and distractor had strong support (top row, Fig. 3c).

To gauge the effectiveness of trial-level modeling, we also analyzed the RT data at the condition level. As typically practiced for condition-level effects in neuroimaging, we aggregated the RT data across trials within each condition through averaging (i.e., complete pooling), thereby assuming the same RT across all trials under each task:

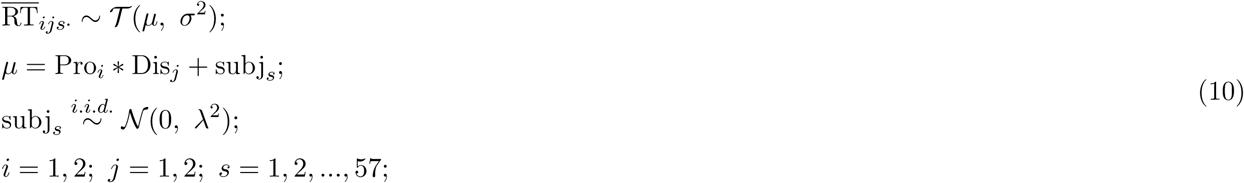

where 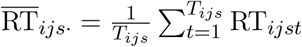. The major difference of the model (10) relative to the trial-level model (9) lies in the omission of terms related to the trial-level effects, trial_*t*_. On the surface, the statistical evidence (Fig. 3d) based on the condition-level model (10) was similar to its trial-level counterpart (Fig. 3c). This is not surprising given the massive evidence for most effects. However, the results also illustrate the higher sensitivity and efficiency of partial pooling relative to complete pooling in that the interaction effect, (`NoRew`−`Rew`):(`Neg`−`Neu`), received only modest support under the condition-based model while being convincingly affirmed by the trial-level model. We note that the interaction was the chief concern in the original study (Padmala et al., 2017), which would not be deemed “statistically significant” under the traditional dichotomous framework. Overall, this example illustrates how data variability is more accurately decomposed and characterized through the trial-level model than the aggregation approach.

#### Neuroimaging data analysis

Time series data were preprocessed using AFNI at each voxel. Steps included cross-slice alignment, cross-TR alignment (mitigation of head motion), cross-subject alignment (normalization to standard space), spatial smoothing (FWHM: 6 mm) and voxel-wise scaling to 100 through dividing the data by the mean signal. To illustrate our modeling framework, we analyzed the data at the ROI level to highlight effective ways to visualize the full results without thresholding, and to demonstrate how BML aids in handling multiplicity. Among the 11 selected ROIs, seven were based on their involvement in attention and executive function more generally: left/right frontal eye fields (FEF), left/right anterior insula (Ins), left/right intraparietal sulcus (IPS), supplementary/pre-supplementary motor area (SMA). We included four additional ROIs, the left/right ventral striatum (VS) and left/right amygdala (Amyg), which are known for their involvement in reward and affective processing, respectively. The ROIs were defined as follows: insula masks were from Faillenot et al. (2017); ventral striatum masks were based on Pauli et al. (2016); amygdala ROIs were defined from Nacewicz et al. (2014); for the remaining regions the peak coordinates of the analysis by Toro et al. (2008) were used to create spherical ROIs.

#### Trial-level effect estimation at the subject level

Trial-level effects were estimated for each subject as follows.^9^ For each ROI, time series data were extracted and averaged across all voxels. The resulting representative time series was analyzed by applying the model (1) with the program 3dREMLfit in AFNI that performs GLS regression combined with REML estimation of the serial correlation parameters in the residuals. Six effects of interest were considered at the condition level: two cue types (`Rew` and `NoRew`) and four tasks (`Rew_Neg`, `Rew_Neu`, `NoRew_Neg` and `NoRew_Neu` factorially combined in terms of the factors Pro and Dis). We compared two approaches: the conventional condition-level method of creating one regressor per condition, and the trial-level approach of modeling each trial with a separate regressor. Each regressor was created by convolving a 1 s rectangular wave with an assumed HRF filter (Gamma variate). Multiple regressors of no interest were also included in the model: separate third-order Legendre polynomials for each run; regressors associated with 6 head-motion effects and their first-order derivatives; and regressors for trials with incorrect responses. In addition, we censored time points for which head motion was deemed substantial (differential movement of 0.3 mm Euclidean distance or above per TR). The 6 runs of data were concatenated with the cross-run gaps properly handled (Chen et al., 2012). With 48 originally planned trials per task, each of the four tasks were modeled at the trial level, resulting in *T*_*ijs*_ (28 ≤ *T*_*ijs*_ ≤ 47) regressors associated with the *i*th prospect and *j*th distractor; each of the two cues were modeled with 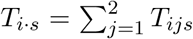 regressors. Each of the error trials and the corresponding cues were modeled separately. For comparison, condition-level effects were also estimated directly for each subject through two approaches. First, each condition was modeled with a regressor that is associated with the *T*_*ijs*_ trials. Second, each condition was modeled with two regressors, one was associated with the average RT across the *T*_*ijs*_ trials while the other captured the modulation effect of RT.

Four approaches were adopted in handling the correlation structure in the residuals: (i) OLS with the assumption of white noise, (ii) AR(1), (iii) AR(2) and (iv) ARMA(1,1), with the latter three models numerically solved through GLS. In addition, we compared the model with AR(2) for the residuals to an AR(2) model with lagged effects of the BOLD signal, as suggested in Westfall et al. (2017). Our comparisons (Appendix C) indicated that AR(2) and ARMA(1,1) for the model residuals rendered similar effect estimates and both slightly outperformed AR(1); thus, all the effect estimates for further analyses were from ARMA(1,1).

Trial-level modeling is vulnerable to the multicollinearity problem. The original experiment was neither intended nor optimally designed for trial-level modeling. Indeed, a few subjects had highly correlated regressors at the trial level between a cue and its subsequent task (correlations among the regressors were below 0.6 for most subjects except for seven who had correlation values above 0.9 among a few regressors). Close inspection revealed that the high correlations were mostly caused by motion effects and the associated data censoring, or by a short separation between a cue and the following task.

Trial-level effects varied substantially without a clear pattern (Figs. 4,13). Across trials, the estimated BOLD response changed substantially, and occasionally showed negative estimates. Such seemingly random fluctuations appeared across all conditions, regions and subjects. Possible factors influencing trial responses include fluctuations in attention, poor modeling and pure noise. Despite the absence of a clear pattern, it is quite revealing to observe some degree of synchronization between the five contralateral region pairs: an association analysis rendered a regression coefficient of 0.73 *±* 0.04 between the region pairs, indicating that a 1% signal change at a right brain region was associated with about 0.73% signal change at its left counterpart (Figs. 4,13).

**Figure 4:**
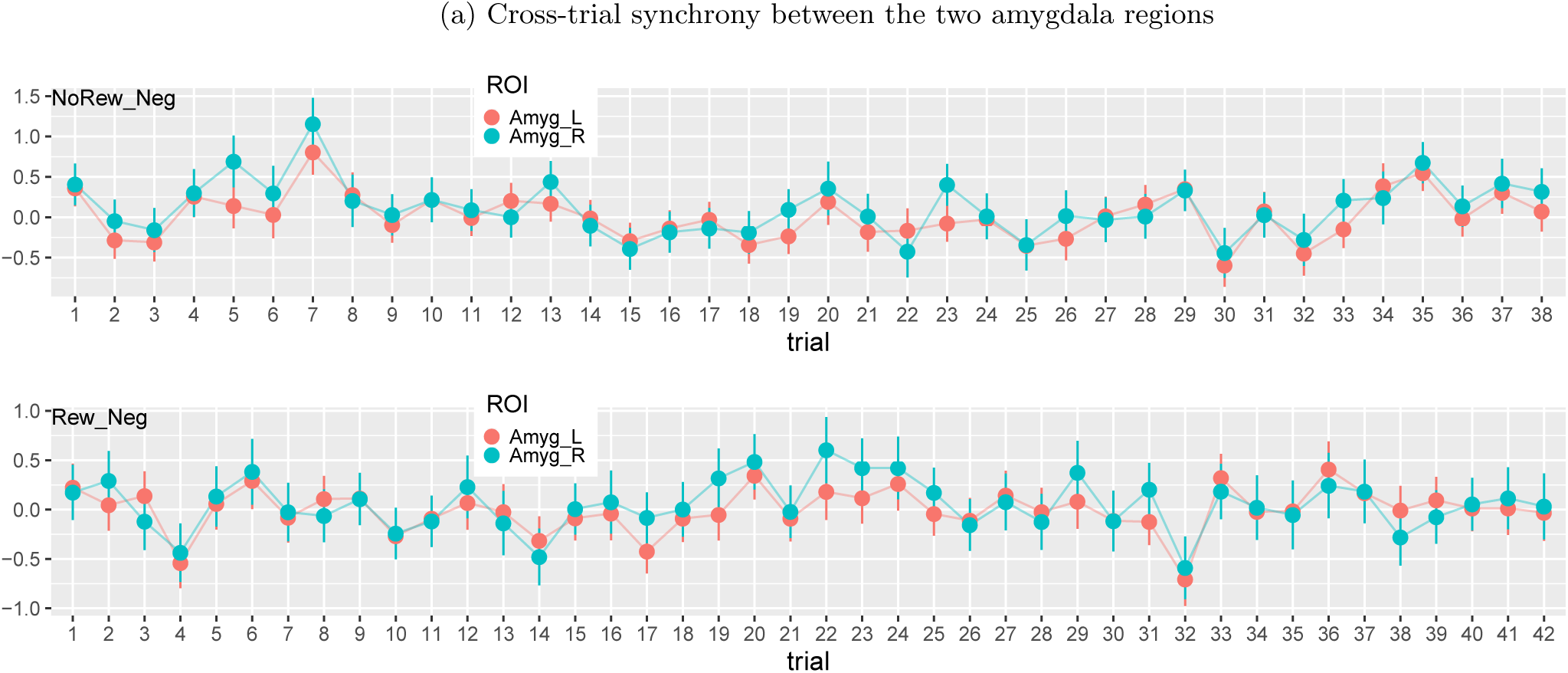
Synchronization among brain regions. The effect estimates (dots) with their standard errors (line segments) were obtained through the GLS model with ARMA(1,1). Some extent of synchrony existed across trials between the left and right amygdalas of a subject under two different tasks of `NoRew_Neg` (upper panel) and `Rew_Neg` (lower panel).

#### Condition effect estimation at the population level

We started with population-level analyses for the four tasks. Inspection of the histogram of effect estimates from the GLS model with ARMA(1,1) in (1) revealed a fraction of outliers that were beyond [-2, 2] in percent signal change (Fig. 5), which were traced mostly to the censoring of time points due to head motion. If not handled properly, extreme values would likely distort the population-level analysis. Unlike the conventional ANOVA framework, BML and LME do not require a balanced data structure without missing data as long as the absence/omission can be considered at random. Specifically, we adopted a BML model with a *t*-distribution to accommodate potential outliers and skewness (Appendix A) and applied to the effect estimates *y*_*ijrst*_ and their variances 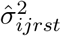 with the indices *i* and *j* coding for the two factors of prospect and distractor:

**Figure 5:**
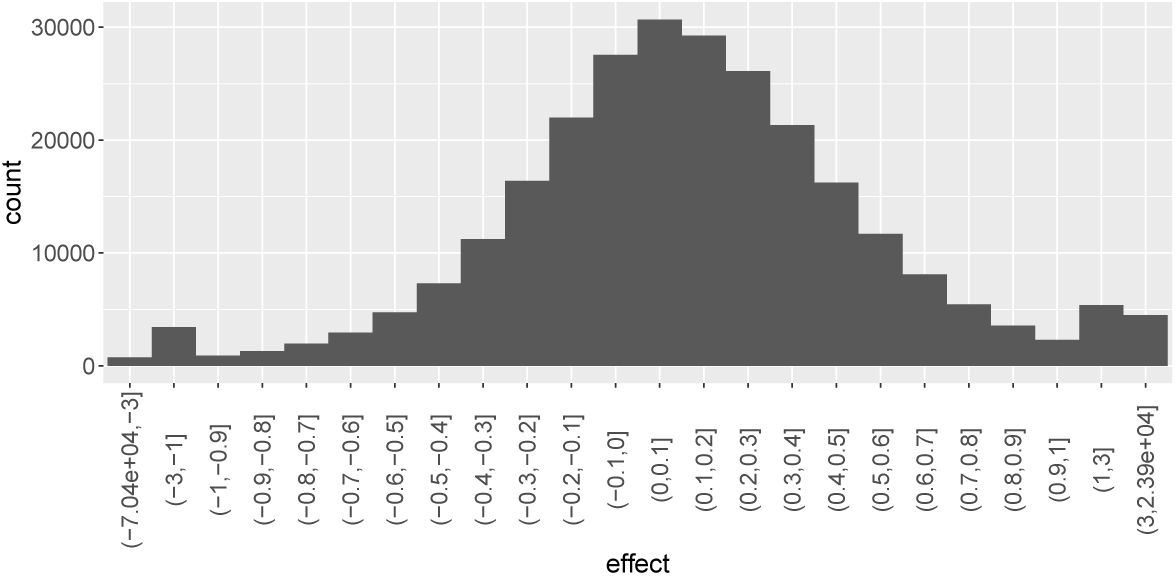
Distribution of the effect estimates from the GLS model with ARMA(1,1). With 11 ROIs and 57 subjects, there were 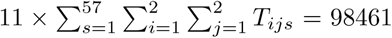 trial-level effect estimates (28 ≤ *T*_*ijs*_ ≤ 47) among the 4 tasks. A small portion (450, 0.42%) were outlying values beyond the range of [-2, 2] with the most extremes reaching -70000 and 23900. To effectively accommodate outliers, the *x*-axis was shrunk beyond (−1, 1).

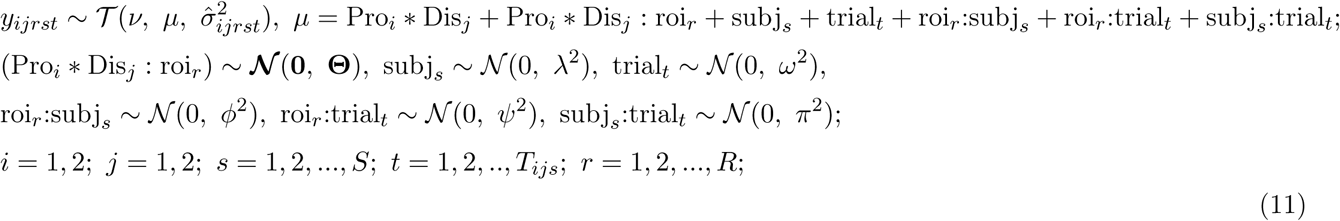

where the 4 × 4 variance-covariance matrix **Θ** captures the cross-region variability among the four tasks. The results are in Fig. 6a.

**Figure 6:**
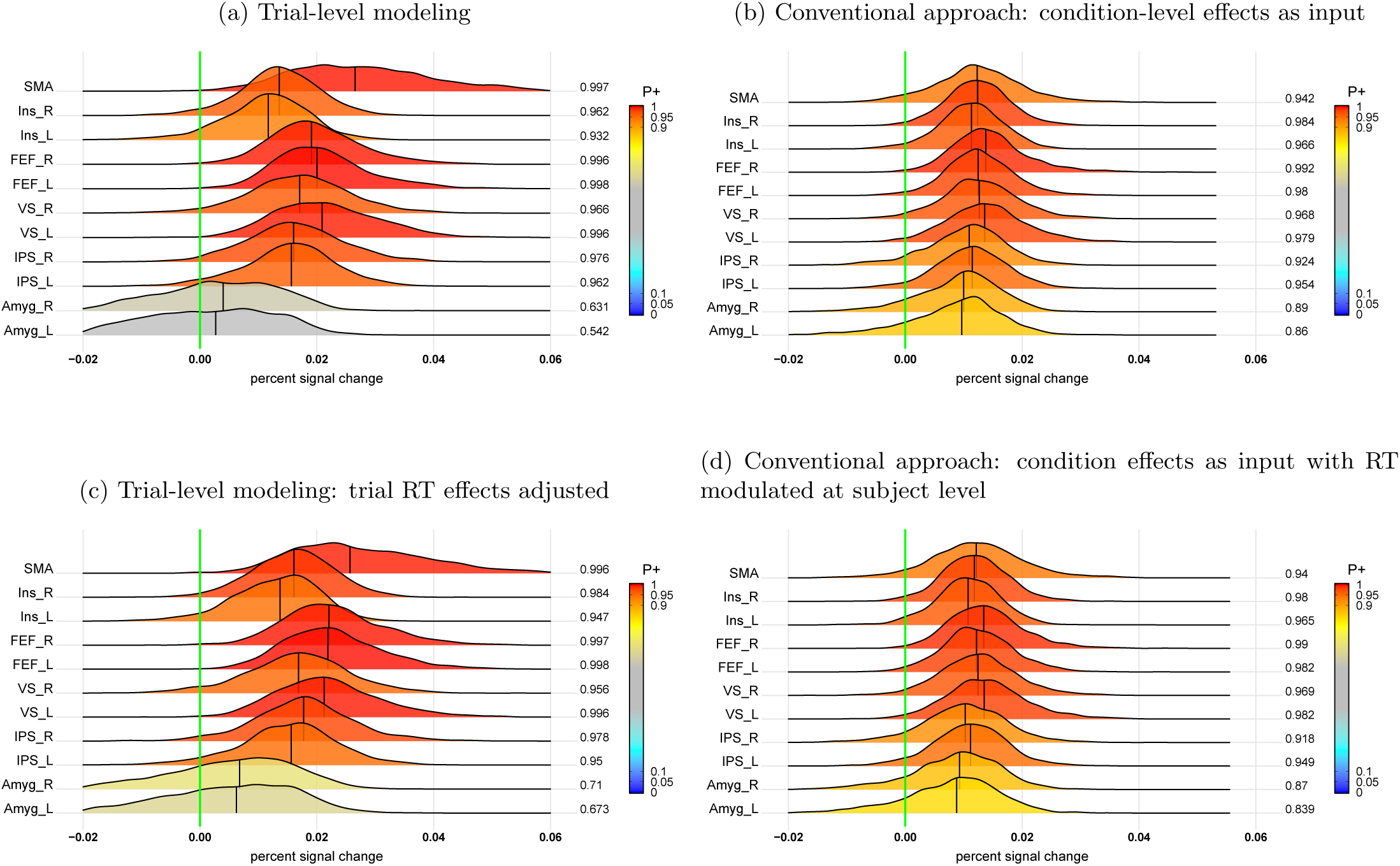
Interaction (`NoRew−Rew`):(`Neg−Neu`) at the population level. The value at the right end of each posterior distribution indicates the posterior probability of the effect being great than 0 (vertical green line), color-coded in the area under each posterior density. Four approaches were adopted to capture the interaction effect: (a) trial-level modeling through the BML model (11); (b) conventional approach: condition-level effects from each subject were fitted in the model (12); (c) covariate modeling: trial-level effects were modeled with RT as a covariate at the population level in the BML model (15); (d) conventional approach: condition-level effects with trial-level RT adjusted from each subject were fitted in the BML model (12).

How do the results in (11) compare to the conventional approach of condition-level modeling through complete pooling? To perform this evaluation, we defined a GLS model (1) with ARMA(1,1) at the subject level that contained six condition-level effects with the assumption that the BOLD response was the same across the trials under each condition. At the population level, task effects *y*_*ijrs*_ and their standard errors 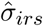 were fitted with the BML model:

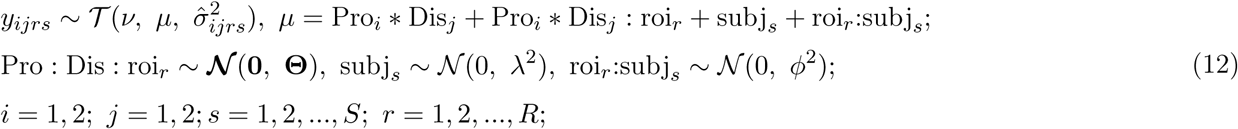

with definitions as before, and where **Θ** is a 4 × 4 variance-covariance matrix for the cross-region variability among the four tasks. Compared to the trial-level modeling approach (Fig. 6a), the condition-level modeling approach produced similar statistical evidence (Fig. 6b), but exhibited inflated reliability (i.e., narrower posteriors), as well as underestimated effect magnitude (densities closer to 0) at most ROIs. In other words, complete pooling tended to homogenize effect estimates and inflate their certainty.

How about the conventional modulation analysis? Under this approach, cross-trial variability is accounted for via a linear modulation of the RT data at the subject level. At the population level we applied model (12) to the condition-level estimates that associated with RT modulation. The resulting interaction effects at the population level (Fig. 6d) were very similar to the ones without the RT modulation (Fig. 6b). Thus, in this case, a modulation regressor did not substantially alter the posterior densities of the interaction effects.

Now we switch to investigate the cue-phase responses with the following BML model applied with the trial-level effect estimates *y*_*irst*_ and standard errors 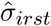 for the cue types `NoRew` and `Rew`:

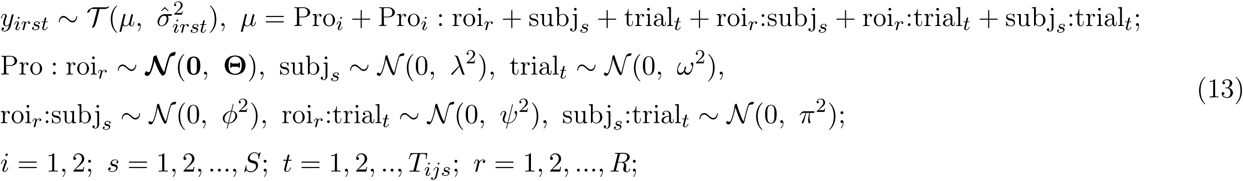

where **Θ** is a 2 × 2 variance-covariance matrix for the cross-region variablity between the two cue types. Most ROIs showed extremely strong evidence for a prospect effect with greater responses during `Rew` relative to `NoRew`(Fig. 7a), although the righ/left amygdala showed weaker support.

**Figure 7:**
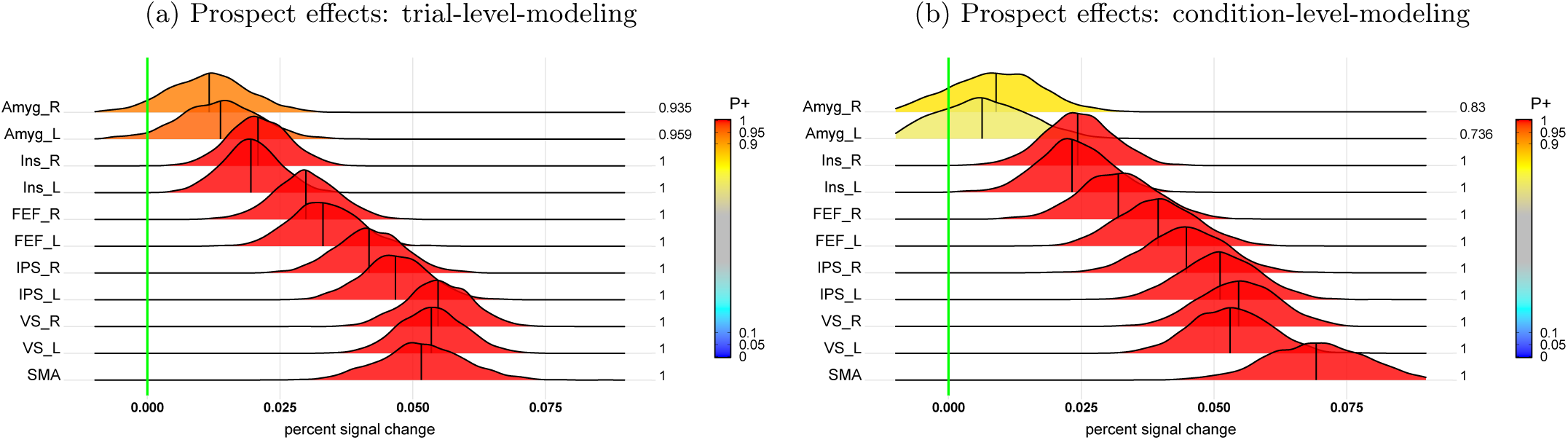
Prospect effect (`Rew−NoRew`) during cue phase at the population level. Even though the two approaches of trial- and condition-level modeling agreed with each other to some extent in terms of statistical evidence for the contrast between `Rew` and `NoRew`, trial-level modeling (a) showed stronger evidence for both left and right amygdala than its condition-level counterpart (b).

Again, to compare the trial-level approach to the conventional condition-level strategy, we fitted with the condition-level effect *y*_*irs*_ and standard errors 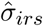 for the cue types `NoRew` and `Rew` with the following:

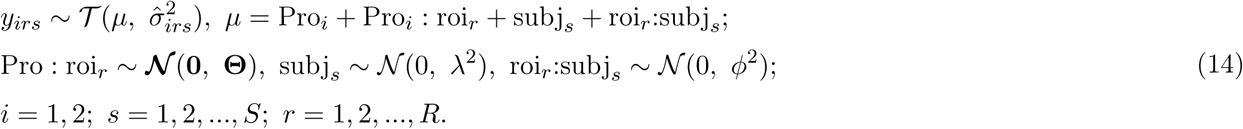

Most of the results from the condition-level approach (Fig. 7b) were similar to the trial-based analysis (Fig. 7a), because of the large effects sizes. However, the condition-level approach did not capture the amygdala effects well, where evidence in their favor was rather weak. In contrast, the trial-based analysis garnered much stronger evidence, and at least the left amygdala would cross a typical one-sided 0.05 statistical threshold (although we believe this dichotomous procedure is detrimental to progress).

#### Association analysis with behavioral data

To probe the linear association between the BOLD response during the task phase and the RT data, we adopted the BML model below for the trial-level effect estimates *y*_*ijrst*_ and their standard errors 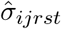:

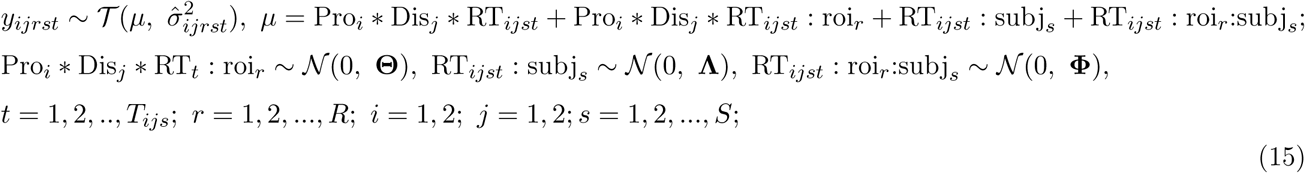

where **Θ, Λ** and **Φ** are 8 × 8, 2 × 2 and 2 × 2 variance-covariance matrices for the respective effects across regions, subjects and the interactions between regions and subjects. As each quantitative variable requires two parameters (intercept and slope) in the model, Pro_*i*_ * Dis_*j*_ * RT_*ijst*_ expands to 8 effects, leading to a 8 × 8 variance-covariance matrix **Θ** for the cross-region variability. There was a strong indication of linearity between the overall task effects and RT in all the ROIs, except in the left and right ventral striatum (Fig. 8a). In addition, when RT was considered as a confounding variable, the interaction (`NoRew`-`Rew`):(`Neg`-`Neu`) showed compatible result (Fig. 6c) as its counterpart (Fig. 6a) from the model (11) without RT modulation.

**Figure 8:**
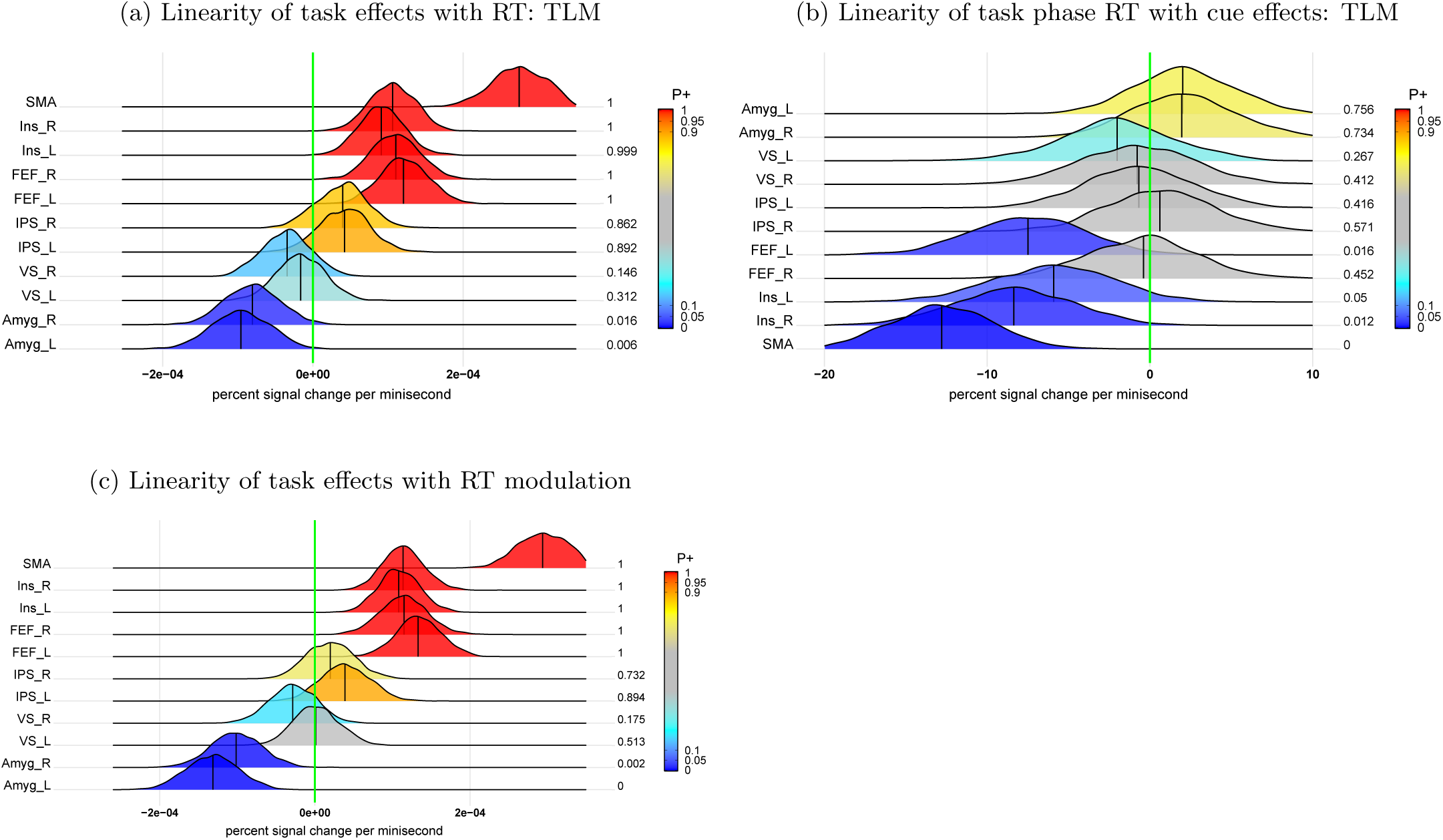
Linear associations of task and cue effects with task phase RT at the population level. (a) Linear association of trial-level effects during the task phase with RT was assessed in the model (15). (b) Linear association of RT during the task phase with the trial-level effects during the cue phase was assessed in the model (16). (c) RT modulation effect during the task phase from the subject level was evaluated in the model (12).

The linear association between cue effects and subsequent task phase RT was also explored. In other words, how were trial-by-trial fluctuations during the cue phase related to task execution? To do so, the following BML model was applied to the behavior data RT_*ist*_ as response variable and the cue effects *x*_*irst*_ as explanatory variable,

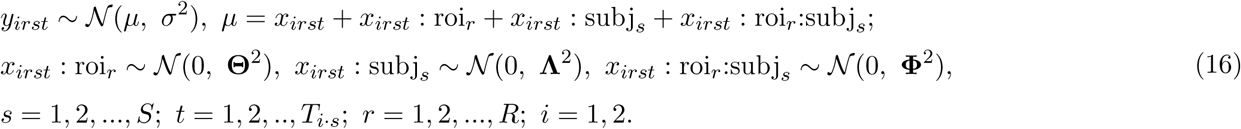

where **Θ, Λ** and **Φ** are 2 × 2 variance-covariance matrices for the respective effects across regions, subjects and the interactions between regions and subjects. Evidence for linear association between the cue phase responses and subsequent behavior was very robust in the SMA, left FEF, and left/right insula (Fig. 8b).

How does the linearity assessed above compare to the conventional modulation nethod? To evaluate such scenario, we applied the formulation (12) to the RT effects *y*_*ijs*_ for the four tasks and their standard error 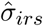 from the modulation analysis at the subject level. Compared to trial-level modeling (Fig. 8a), the RT effects based on modulation (Fig. 8c) showed very similar results.

Is linearity too strong an assumption even though frequently assumed in investigating the relationship between FMRI signals and covariates of interest (e.g., RT)? To address this question, we focused on one of the cue/task combinations, namely `Rew_Neg`, involving reward cues and negative distractors. The trial-level effects *y*_*rst*_ and its variance 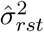 under the task `Rew_Neg` were fitted as follows,

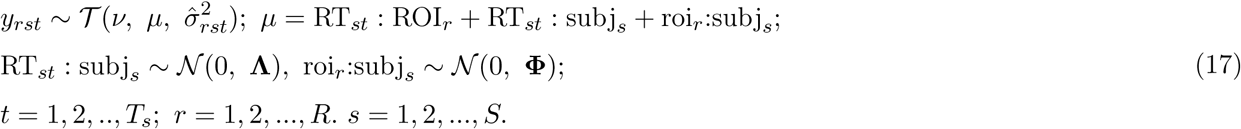

Separately, a nonlinear function was applied to the RT via smoothing splines:

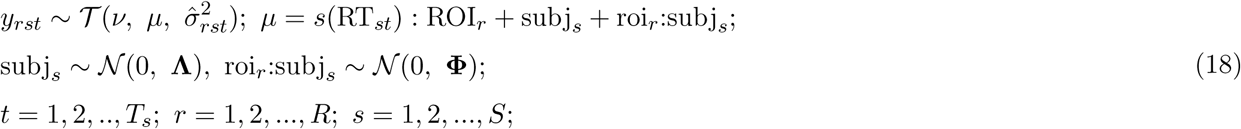

where the smoothing function *s*(*·*) adopts a cubic spline basis defined by a set of knots spread evenly across the RT range and penalized by the conventional integrated square second-derivative cubic-spline term. With the narrow range of the RT values, the dimensionality of the basis expansion (i.e., number of knots) was set to 10 through a generalized additive model so that simplicity was balanced against explanatory power. Due to the adaptive nature of regularization, the exact choice of knots is not generally critical as long as it is large enough to represent the underlying mechanism reasonably well, but small enough to maintain adequate computational efficiency (Wood, 2017).

Fitting the data with linear and nonlinear models yielded some similarities, but multiple differences were also observed (Fig. 9). For example, linear fitting revealed positive trends at the SMA and the right/left IPS between task responses and the corresponding RT (Fig. 9a), whereas the spline fittings uncovered more complex relationships (Fig. 9b). The largely parallel trends observed across the contralateral region pairs provide some validation for both the linear and nonlinear fittings. Nevertheless, the nonlinear results suggest that linearity is likely too strong an assumption across the whole RT range; thus, the statistical evidence for linearity might have been inflated. For example, linearity might be applicable for certain RT ranges, but support for it might be limited at lower and higher values of RT with fewer data points. In addition, the uncertainty under the model assuming linearity (17) appears to have been considerably underestimated, especially when RT is away from the central values.

**Figure 9:**
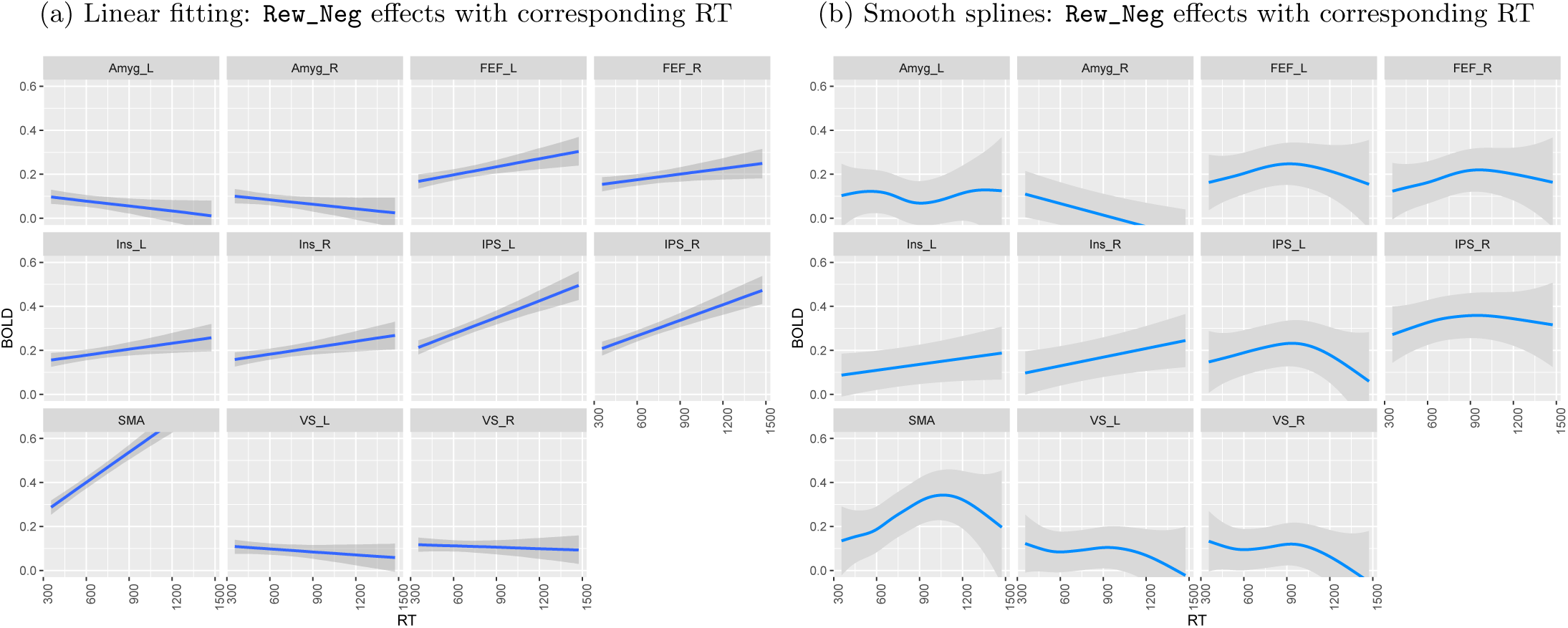
Comparisons of association analysis under the task `Rew_Neg` between linear fitting and smoothing splines. For better visualization on the dependence of trial-level effects on RT, the trends are shown with their 95% uncertainty bands. (a) Linear fitting was assessed in the model (17). (b) Association analysis was evaluated through smoothing splines in the model (18).

**Figure 10:**
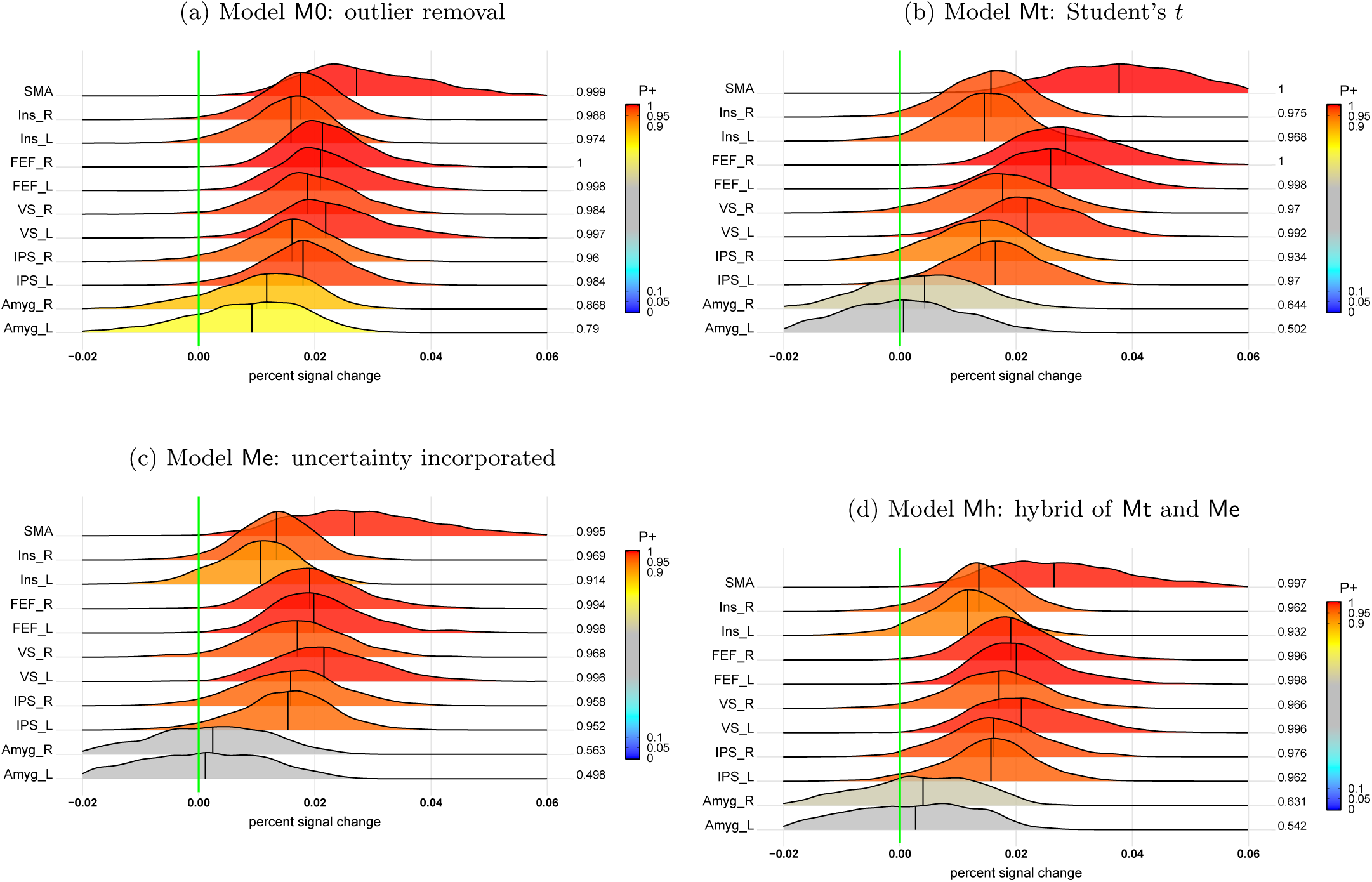
Interaction (`NoRew−Rew`):(`Neg−Neu`) at the population level through four different BML versions. The value at the right end of each line indicates the posterior probability of the effect being great than 0 (vertical green line), color-coded in the area under each posterior density. Four BML models were adopted to handle outliers: (a) M0: brute force removal of values outside [-2, 2]; (b) Me: incorporation of uncertainty for effect estimates; (c) Mt: adoption of *t*-distribution to accommodate outliers and skewness; and (d) Mh: hybrid of Me and Mt with both the uncertainty of effect estimates and *t*-distribution.

## Discussion

Experimental sciences aim for generalizability. In doing so, they draw inferences about populations – idealized, theoretical constructs – from samples of, for example, trials and subjects. The ability to generalize is typically achieved through framing the cross-sample variability with an appropriate statistical distribution (e.g., Gaussian, Student-*t*). At the same time, when carefully choosing a stimulus set and subjects, an experimentalist obviously aims to draw conclusions that reach beyond the particular instances utilized. As recognized by Westfall et al. (2017), the standard analysis framework in neuroimaging does not lend itself to such a goal in the case of stimuli, and the same logic employed in bridging subjects to “population effects” is required. The present paper develops a two-level approach that is well adapted to the current analytical streamline; in addition, partial pooling is applied to trial-level effects at the population level to tackle the generalizability problem. We illustrated the effectiveness of the approach via a series of analyses of an FMRI dataset from a rich experimental paradigm with multi-phase trials, including cue stimuli and subsequent task execution.

### Why should we more accurately account for trial-level variability?

The issue of cross-sample variability has been recognized for several decades across multiple research areas. For example, Clark (1973) pointed out that neglecting the problem of the “language-as-fixed-effect fallacy … can lead to serious error” (p. 335), and even alluded to earlier warnings that were largely ignored in the literature, including the report by Coleman (1964). Given the common practice of aggregation across trials, even classical experiments such as Stroop and flanker tasks, which one would anticipate to show high reliability, yielded lackluster results (Rouder and Haaf, 2019).

In neuroimaging, trial-level variability is typically bundled together and flattened with the residuals of the GLS model at the subject level when condition-level effects are of interest. Such practice assumes that all trials have exactly the same BOLD response. In our analysis of an FMRI dataset, considerable variability was observed across trials (Figs. 4, 13). Although the attribution of this variability to “pure noise” cannot be excluded, our results collectively point in a different direction: the variability is meaningful. Indeed, we interpret our results as suggesting that, without directly capturing trial-level effects, population-level estimates can be compromised. In particular, our results suggest that condition-level modeling may underestimate effect magnitudes, while in some cases overestimating certainty (Fig. 6d,e; Fig. 7).

Ignoring cross-trial variability also leads to the loss of legitimately being able to generalize beyond the stimulus set used. As emphasized by Clark (1973), a serious implication is that studies will be “particularly vulnerable to lack of replicability”. Generalization from a stimulus set to a category requires proper model construction (Coleman, 1964; Clark, 1973; Westfall et al., 2017; Baayen et al., 2008; Yarkoni, 2019). When condition-level effects are inferred without accounting for cross-trial variability, technically speaking, the conclusions are applicable only to the particular trials in the experimental design, not even to similar cases from the same category. Trials are typically conceptualized as originating from a population that follows specific distributional assumptions (e.g., Gaussian), thus supporting the generalization from specific trials to the associated category. In addition, the explicit accountability of cross-trial variability provides more accurate characterization of both the effect magnitude and its uncertainty. Currently, modeling cross-subject variability is considered standard in the field as a way to draw population-level inferences. We believe the same should be considered for cross-trial variability.

Modeling cross-trial variability is important even if the difference in statistical evidence is small practically. First, although the fact that cross-trial variability diminishes as the sample size increases (see expression (4)), it is not practically possible to realistically determine the “required” number of trials. As the sample size in our experimental data was reasonably large (57 subjects and 28-47 trials per task), the differences in statistical evidence between trial- and condition-level modeling were not large (e.g., Fig. 6a,b). Nevertheless, meaningful differences were observed, such as the strength of the evidence for cue effects in the amygdala (Fig. 7). More importantly, condition-level modeling showed distortions (e.g., underestimation) in both the magnitude and uncertainty of effect estimates (e.g., Fig. 6a,b). Furthermore, whereas investigators are generally cognizant of the need to employ enough subjects, awareness about requirements about trial sample size remains limited. We believe that considerations about trial sample size should be on a comparable footing as those of the number of participants.

Modeling trial-level responses is also of great value when studying brain-behavior relationships. A rich literature has investigated how trial-by-trial fluctuations in behavior are associated with intertrial variability (Ress et al., 2000; Pessoa et al., 2002; Pessoa and Padmala, 2005; Sapir et al., 2005; Lim et al., 2009). Many of these studies have proposed that the most likely source of the association is related to trial-by-trial changes in attention. Another potential source (Fox et al., 2006; Fox et al., 2007) is that intrinsic signal fluctuations account for much of intertrial variability in human behavior. Specifically, spontaneous fluctuations of the BOLD signal in resting-state studies also contributed to fluctuations during behavioral tasks. Evidences showed that ongoing intrinsic activity accounted for 60% of the variability in brain responses during a simple button-pressing task (Fox et al., 2007). Overall, although cross-trial variability has been framed in terms of issues of “fixed” versus “random” effects (Westfall et al., 2017), we believe it is of potential value to conceptualize the problem from a much broader perspective.

We also investigated the performance of the integrative LME modeling approach previously proposed by Westfall et al. (2017), which utilizes a different modeling approach than our proposed one, from the following perspectives.

#### 1) AR structure

Their model aims to explicitly accounts for the serial correlation of the times series with lagged effects as explanatory variables, instead of capturing the AR structure in the residuals as typically practiced in the field. Such an approach remains controversial (e.g., Achen, 2000; Keele and Kelly, 2006; Bellemare et al., 2017; Wilkins, 2018), and it had a dramatically large impact on the results with the present dataset (Figs. 13b,14 in Appendix C). In addition, the following assumptions of their model are likely inaccurate: that all subjects share the exactly same AR structure, as well as the same baseline and cross-trial effects.

#### 2) Assumption of white noise

Due to the violation of endogeneity (e.g., omitted variables, measurement error), the residuals in their integrative LME model (7) would be correlated with the lagged response variables. Accordingly, it would still be important to model the temporal structure in the residuals.

#### 3) Focus on statistical evidence

Westfall et al. (2017) reported statistic values without accompanying effect estimates. We believe this practice, which is common in neuroimaging, is problematic (see Chen et al., 2017) because it leads to information loss by reducing the effect estimates to a simple binary statement. Instead, we advocate in favor of documenting voxel- or region-wise magnitudes and their respective uncertainty; in addition, study reports without revealing effect magnitudes contribute to the reproducibility problem.

At the population level, what are the practical consequences of ignoring cross-trial variability? Westfall et al. (2017) made the strong cautionary warning that it could produce inflation 1.5 to 3 times of the values of the relevant statistic employed. The changes that we observed, important as they were, were less dramatic, and in some cases involved *deflation* of statistical evidence (e.g., cue effects, Fig. 7). In addition, some substantial differences in effect magnitude and reliability were observed too (e.g., interactions between prospect and distractor, Fig. 6a,b). It is possible that differences between the present approach and that by Westfall et al. (2017) were partly influenced by some modeling choices including AR handling and other modeling assumptions. In this context, it is informative that we also observed a small amount of deflation in statistical evidence when data were aggregated across trials for the behavioral data (Fig. 3). In fact, our observations are more aligned with a similar assessment of test-retest reliability in psychometrics (Rouder and Haaf, 2019). We emphasize that our observations here were based on a relatively large number of trials, and we recognize the difficulty of making a general assessment about the practical impact due to the involvement of multiple factors, such as task type, sample size, the number of trial repetitions, and the type of experimental design (event-related vs block). The risk of statistical evidence inflation, when cross-trial variability is not properly accounted for, can be substantial when trial samples are not large, as demonstrated by Westfall et al. (2017).

### Benefits of the two-level modeling approach

We propose a unified statistical platform that addresses the generalizeability issue through a two-level modeling approach (Table 1). Instead of adopting the conventional *complete pooling* (all trials essentially averaged), we directly estimate the trial-level effects at the subject level through *no pooling* (parameter estimates obtained for each trial separately). At the population level *partial pooling* is adopted via a hierarchical model to achieve generalizability from specific trials to condition category. Note that although we adopted a Bayesian platform here, the framework can also be implemented with an LME model, if one employs a whole-brain voxel-wise approach and is not interested in partial pooling across ROIs. We now discuss a few strengths of our approach.

**Table 1:** Parallelism between cross-subject and cross-trial variability

| Strategy | No Pooling | Partial Pooling | Complete Pooling | Association |
| --- | --- | --- | --- | --- |
| **Info. Sharing** | none (individual) | some (adaptive) | full (uniform) | some (associative) |
| **Subject Effect** | unique for $s$th subject:<br>$\xi_s \sim \mathcal{U}(-\infty, \infty)$<br>(**current practice**) | regularized:<br>$\xi_s \sim \mathcal{N}(0, \lambda^2)$<br>(**current practice**) | same across subjects:<br>$\xi_s = 0$<br>(‘**fixed effects**’) | subject-level<br>covariate<br>(e.g., age) |
| **Trial Effect** | unique for $t$th trial:<br>$\eta_t \sim \mathcal{U}(-\infty, \infty)$<br>(**proposed practice**) | regularized:<br>$\eta_t \sim \mathcal{N}(0, \omega^2)$<br>(**proposed practice**) | same across trials:<br>$\eta_t = 0$<br>(**current practice**) | trial-level<br>behavior<br>(e.g., RT) |
| **Variance Prior** | ∞ | finite ($\lambda^2, \omega^2$) | 0 | - |
| **Properties** | unbiased but unreliable | biased but more accurate | homogenized | - |
| **Applicability** | prep. for population<br>analysis; predictions;<br>cross-region correlations | subjects to population,<br>trials to category | - | controllability,<br>correlation |

#### 1) Computational feasibility

Our two-level approach attains computational tractability by segregating subject and population analyses. Importantly, at the same time, with subject-level uncertainty (standard errors) carried to the population level, any potential information loss is likely minimal relative to a “single-step” integrative method. The BML models can be implemented through the R package brms, which builds on top of the Stan language. At the same time, equivalent LME models can be performed at the voxel level through the program 3dLMEr publicly available in the AFNI suite.

#### 2) Flexibility and adaptivity

The two-level approach can be adopted for several research objectives, including: (a) condition-level effect estimation at the population level; (b) classification and machine learning applications utilizing trial-level estimates; (c) correlativity analysis based on trial-level effects; and (d) brain-behavior association. At a basic level, close examination and visualization of the trial-level effects become possible (e.g., synchrony between contralateral regions or among regions in a network).

#### 3) Outlier handling

The possibility of outliers and data skew needs to be carefully considered in trial-by-trial analyses. The present framework flexibility accounts for these possibilities by (a) incorporating reliability information and (b) regularizing the estimation via, for example, Student’s *t*-distribution.

#### 4) Modeling options

Standard modulation analysis is able to investigate associations between behavioral variables and BOLD responses at the trial level. Our approach allows the evaluation of brain-behavior associations by assuming a linear relationship at the population level. In addition, the approach offers the investigator the opportunity to flexibly explore nonlinear relationships with smoothing splines.

### Additional trial-level modeling issues

Our study also sheds insights about other aspects of FMRI data analysis.

#### 1) The importance of modeling AR structure in the residuals

Our investigation on AR effects (Appendix C) confirmed a previous study (Olszowy et al., 2019) and indicated that the AR structure in the residuals varies substantially across regions, tasks and subjects. Therefore, we recommend that, to obtain reasonably accurate standard errors for effect estimates, a GLS model with the temporal structure in the residuals be accounted for with preferably AR(2) or ARMA(1, 1) for a TR around 2 s. With shorter TRs, a higher-order AR structure would be likely needed (Olszowy et al., 2019; Luo et al., 2020).

#### 2) Incorporating effect uncertainty in population analysis

Should standard errors of effect estimates be modeled at the population level? Previous studies suggested that the benefit was minimal (Mumford et al., 2009; Chen et al., 2012; Olszowy et al., 2019). However, since trial-level modeling is more prone to multicollinearity and may result in unreliable effect estimates, uncertainty information provides a robust mechanism to counter the impact of outliers at the population level. As the accountability of the serial correlation in the residuals of the time series regression model is influential on the accuracy of the standard error for each effect estimate, it becomes important to more accurately model the AR structure at the subject level.

#### 3) Multicollinearity

As each trial is estimated as a separate regressor, careful experimental design and trial-order optimization should be considered, especially when the TR is relatively short. In particular, stimulus timing can be determined to reduce multicollinearity using tools such as RSFgen in AFNI or optseq^10^. For analyses based on behavioral performance which cannot be optimized in advance, particular attention should be paid to multicollinearty. However, given the overall two-stage estimation procedure, mutlicollnearity may pose less of a problem than typically assumed, although it will likely affect the precision of the estimated effects.

#### 4) Hemodynamic response modeling (HDR)

The HDR can vary in shape in several ways, including response delay and speed, peak width, recovery length, and presence of onset and recovery undershoots. The HDR variability may occur across trials, tasks, brain regions, subjects and groups. It might be economical and efficient to adopt a generic and fixed-shape HDR function if the model provides a reasonably close fit to the data, but such a methodology likely becomes overly simplistic to model the HDR across more general scenarios. Importantly, it is possible that some of the cross-trial variability is due to the poor HDR fitting currently adopted in the field. At present, two approaches are available to provide fitting flexibility. One is to include one or two more adjusting functions (e.g., temporal derivative and dispersion curve), and the other is to adopt a data-driven approach through response estimation with multiple basis functions (e.g., linear or cubic splines). However, due to the large number of regressors involved, trial-level modeling is largely confined to using a fixed-shape HDR, which limits the ability of the approach to handle deviations from more canonical-shaped responses.

## Conclusions

In the present study, we investigated the extent and impact of trial-by-trial responses in FMRI data. While the importance of trial-leveling modeling has been raised previously, we propose and evaluate several new modeling strategies to address the issue. Using real FMRI data, we demonstrated the benefits of these new approaches in terms of both mathematical structure and in terms of interpretable outcomes. At the trial level, responses were estimated through a GLS model with serial correlations accounted for, whereas population-level analysis was carried via a hierarchical model that effectively characterized effect structure, allowing generalizability from the specific stimuli employed to the generic category. Additional applications of the approach employed here include the analysis of brain-behavior associations, trial-based correlation analysis, as well as trial-level classification and machine learning.

## Acknowledgments

The research and writing of the paper were supported (GC, PAT, and RWC) by the NIMH and NINDS Intramural Research Programs (ZICMH002888) of the NIH/HHS, USA. LP’s research was supported in part by NIMH (R01 MH071589 and R01 MH112517). We are appreciative of the technical support from the Stan (Carpenter et al., 2017) and R (R Core Team, 2019) communities. Most of the modeling work was performed in Stan through the R package brms (Bürkner, 2018), and the figures were generated with the R package ggplot2 (Wickham, 2009). Kelly Morrow, Chirag Limbachia and Anastasiia Khibovska provided assistance in generating some of the figures, and Kelly Morrow for help in creating the regions of interest. We thank the anonymous reviewers for their critical reading with thoughtful suggestions that helped improve and clarify our manuscript.

## Appendix A

### Four BML models for trial-level modeling

Four different BML models are considered for trial-level modeling. Following our recent Bayesian approach (Chen et al., 2019a), we formulate the models within a single integrative platform at the level of region of interest (ROI) to capture the hierarchical structure among three intersecting levels: subjects, trials and regions. Specifically, the trial-level effect estimates *y*_*str*_ are modeled as follows:

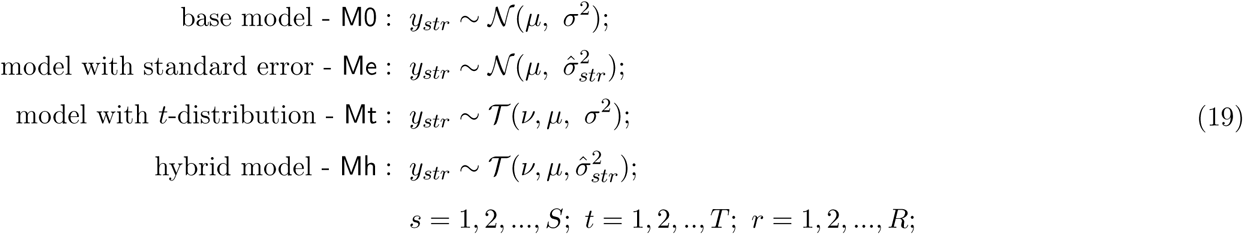

where *r, s* and *t* index the ROIs, subjects and trials, and the parameter *ν* is the number of degrees of freedom for the *t*-distribution. The four BML models differ along two dimensions in a crossed manner: (a) whether the uncertainty information 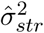 (effect variance from the subject level) is incorporated and propagated from the subject to population level, and (b) whether Gaussian *𝒩* or Student’s *t*-distribution *𝒯* is assumed for the response variable. Under the Bayesian framework, the BML models (19) are expressed as a distribution or likelihood function, rather than as an equation (like the LME model in (2)). Hence, the parameter *µ* in the four models (19) can be further specified as follows:

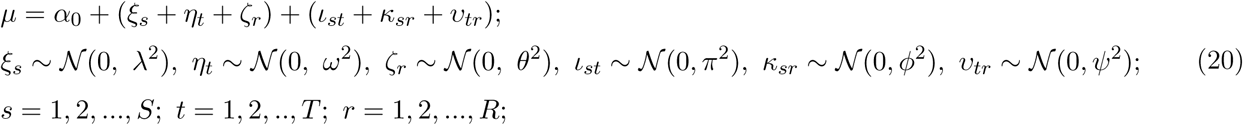

where the indices *s* and *t* code for subjects and trials, respectively; *α*_0_ is the intercept; *ξ*_*s*_ and *η*_*t*_ represent cross-subject and cross-trial effects; *ζ*_*r*_ accounts for the effect of the *r*th region. Compared to its LME counterpart (2), the four BML models incorporate brain regions (indexed by *r*), augmenting the LME model to a platform with three crisscross levels. Due to the addition of this cross-region dimension, it is possible to further include the three two-way interaction terms, *ι*_*st*_, *κ*_*sr*_ and *υ*_*tr*_ among the three effects of subjects, trials and regions (with parentheses grouping the three single levels and their interactions).

The relationships among the four BML models are as follows. Relative to the base model M0, Me takes into account the precision of the effect estimate *y*_*str*_ by utilizing the standard error 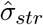 from the subject level. To handle outliers and skew in the data *y*_*str*_, we replace the Gaussian distribution with Student’s *t*-distribution and formulate the third BML model, Mt. Thanks to its leptokurtic property (i.e., having “heavy tails”), the *t*-distribution (with the Gaussian distribution as its asymptote) has increasingly more mass in the tails as the degrees of freedom decrease, effectively counteracting the potential impact of outlying values. Lastly, the models Me and Mt can be combined to incorporate uncertainty and to handle outliers simultaneously, leading to the hybrid Mh. In all model versions, the intercept *α*_0_ can be expanded with terms to accommodate terms such as subject-grouping and quantitative covariates.

The four BML models generated relatively similar posterior distributions (Fig.10a-d). First, the base model M0 overestimated the effect magnitude for the two amygdala ROIs. To appreciate the differences between the four models, consider the degrees of freedom for the *t*-distribution, *ν*, in Mt which is adaptively estimated from the shape of the data distribution. The strength of *t*-distribution in handling heavier tails and potential outliers is demonstrated by the small degrees of freedom estimated as *ν* = 3.24 *±* 0.03 in Mt (cf. the Gaussian distribution of M0 with *ν* = ∞). The inclusion of uncertainty in Me also allowed effective handling of extreme values (similar estimates as Mt, in particular for the left/right amygdala), noticeable in the decreased role of the *t*-distribution in the hybrid model Mh, where *ν* = 61.8 *±* 5.1. Overall, instead of relying on a predetermined threshold value to handle outliers in M0, models Mt, Me and Mh offer principled approaches to adjusting to the shape of the data distribution and the presence of potential outliers. The important role of standard errors in the models Me and Mh necessitates the accurate accountability of the serial correlation in the GLS model (1). In the main text of the paper, the model Mh is adopted, whenever applicable, for its more adaptive formulation.

## Appendix B

### Hyperpriors adopted for BML modeling

The prior distribution for all the lower-level (e.g., trial, ROI, subject) effects considered here is Gaussian, as specified in the respective model; for example, see the distribution assumptions in the BML model (20). If justified, one could adopt other priors like Student’s *t* for the effects across trials, regions and subjects, just as for the likelihood (or the prior for the response variable *y* in the BML model (19)). In addition, prior distributions (usually called hyperpriors) are needed for three types of model parameters in each model: (a) population effects or location parameters (“fixed effects” under LME, such as intercept and slopes), (b) standard deviations or scaling parameters for lower-level effects (“random effects” under LME), and (c) various parameters such as the covariances in a variance-covariance matrix and the degrees of freedom in Student’s *t*-distribution. Noninformative hyperpriors are adopted for population effects (e.g., population-level intercept and slopes). In contrast, weakly-informative priors are utilized for standard deviations of lower-level parameters such as varying slope, subject-, trial- and region-level effects, and such hyperpriors include a Student’s half-*t*(3, 0, 1) or a half-Gaussian *N*_+_(0, 1) (a Gaussian distribution with restriction to the positive side of the respective distribution).

For variance-covariance matrices, the LKJ correlation prior (Lewandowski, Kurowicka, and Joe, 2009) is used with the shape parameter taking the value of 1 (i.e., jointly uniform over all correlation matrices of the respective dimension). Lastly, the standard deviation *σ* for the residuals utilizes a half Cauchy prior with a scale parameter depending on the standard deviation of the input data. The hyperprior for the degrees of freedom, *ν*, of the Student’s *t*-distribution is Γ(2, 0.1). The consistency and full convergence of the Markov chains were confirmed through the split statistic 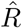 being less than 1.1 (Gelman et al., 2013). The effective sample size (or the number of independent draws) from the posterior distributions based on Markov chain Monte Carlo simulations was more than 200 so that the quantile (or compatibility) intervals of the posterior distributions could be estimated with reasonable accuracy.

## Appendix C

### Handling autocorrelation in FMRI data

The amount of temporal correlation embedded in the residuals of the time series regression with trial-level modeling was substantial with large variations across regions, tasks and subjects (Fig. 11). Specifically, the overall serial correlation across the 11 ROIs and 57 subjects was 0.50 *±*0.20, 0.47 *±*0.28 and 0.33 *±*0.38 assessed from the AR(1), AR(2) and ARMA(1,1) models, respectively), indicating that some large amount of effects were not properly accounted for through the explanatory variables. With condition-level modeling, cross-trial fluctuations would become part of the residuals; thus, the AR effects would be different and likely stronger, The performances of the OLS approach were compromised due to the presence of persistent temporal correlation in the model residuals. Based on the Gauss-Markov theorem, the OLS method would still provide consistently unbiased estimates, with the caveat that the precision for the effect estimates tends to be inflated. However, the asymptotic property of the unbiasedness heavily relies on a large sample size, which cannot necessarily be met nor easily predetermined in real practice. With the current dataset, the OLS solutions showed some extent of over- and under-estimation compared to the three AR models (Fig. 12). In addition, a slight amount of underestimated uncertainty (or inflated precision) about the OLS effect estimates is evident compared to their AR counterparts (Fig. 12).

**Figure 11:**
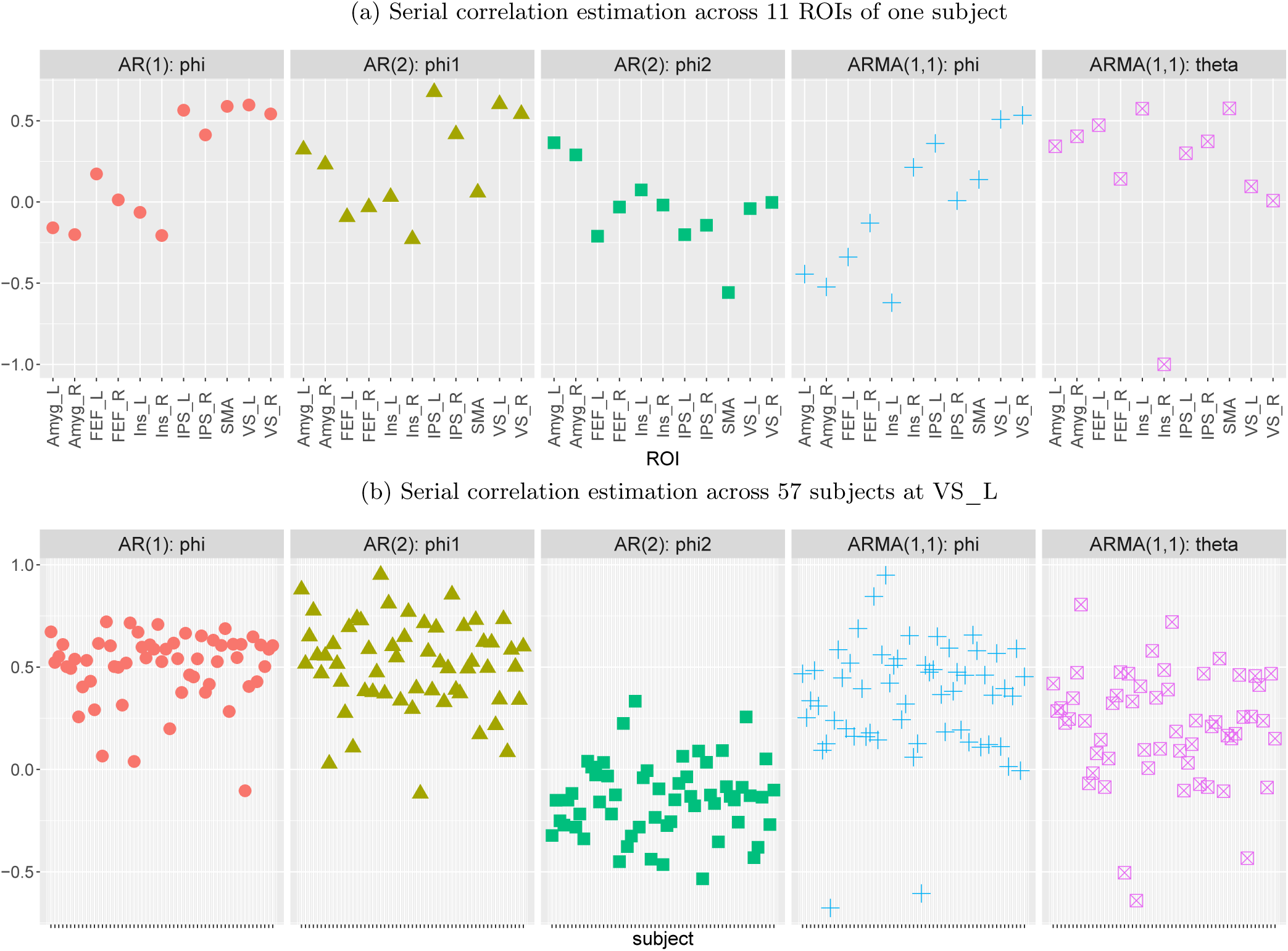
Variations of temporal correlation across regions and subjects. The overall average first-order AR parameter of trial-level modeling across all the 11 ROIs and 57 subjects was 0.50 0.20, 0.47 0.28 and 0.33 0.38 for AR(1), AR(2) and ARMA(1,1), respectively; the second-order parameter for AR(2) and moving average parameter for ARMA(1,1) were − ∓ 0.13 −0.17 and 0.18 ± 0.34, respectively. The relative magnitude of these AR parameters indicated that the first AR parameter captured substantially large proportion of the serial correlation while the second parameter in AR(2) and ARMA(1,1) remained helpful.

**Figure 12:**
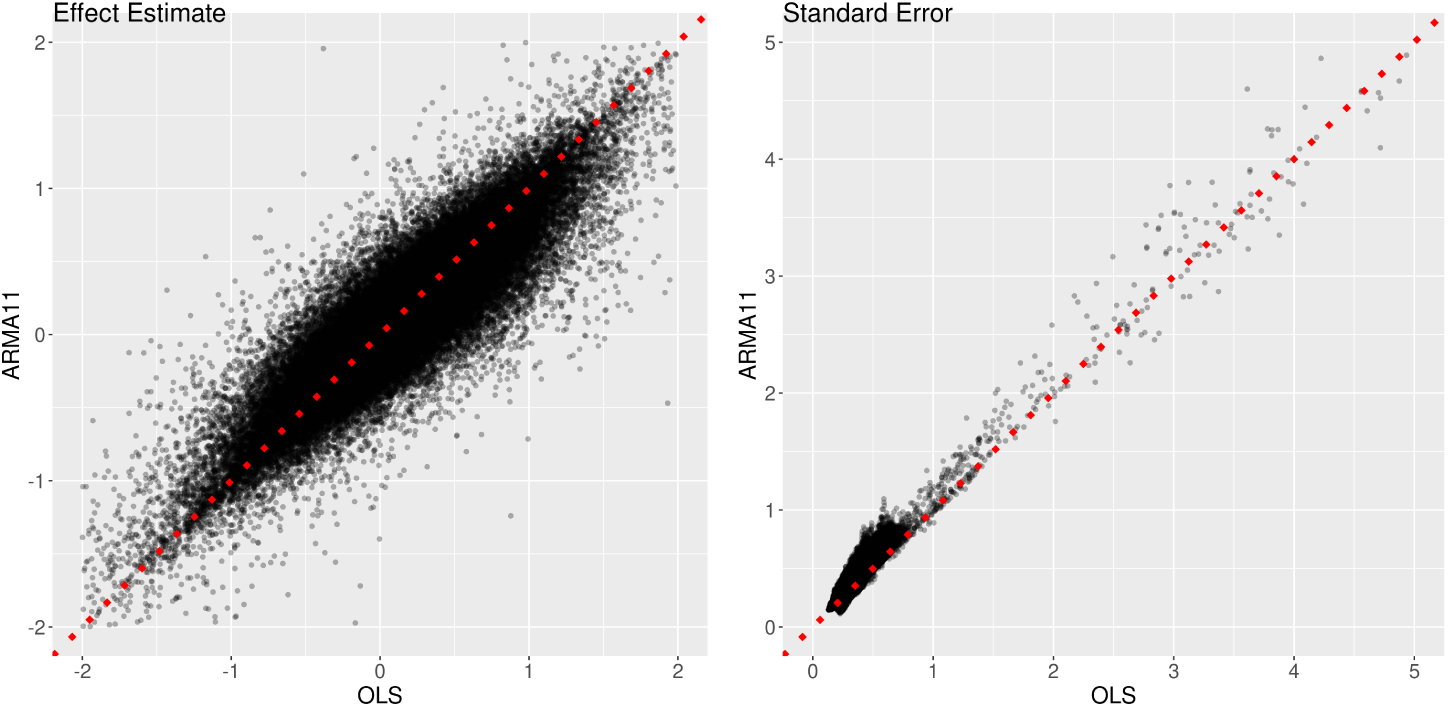
Comparisons of OLS and ARMA(1,1) in effect estimate and uncertainty. The effect estimates (left) and their standard errors (right) are shown for the total 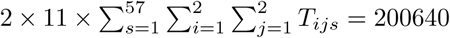 trial-level effects among the two cues and four tasks. The theoretical unbiasedness of OLS estimates can be verified by the roughly equal number of data points on the two sides of the diagonal line (dotted red). However, the instability of OLS estimation is shown by the fat cloud surrounding the diagonal line: slightly overestimation (or underestimation) of OLS was shown by 52.7% (or 45.5%) of data points above (or below) the *x*-axis. The precision inflation of OLS can be assessed by the proportion of data points (97.5%) above the dotted red line.

**Figure 13:**
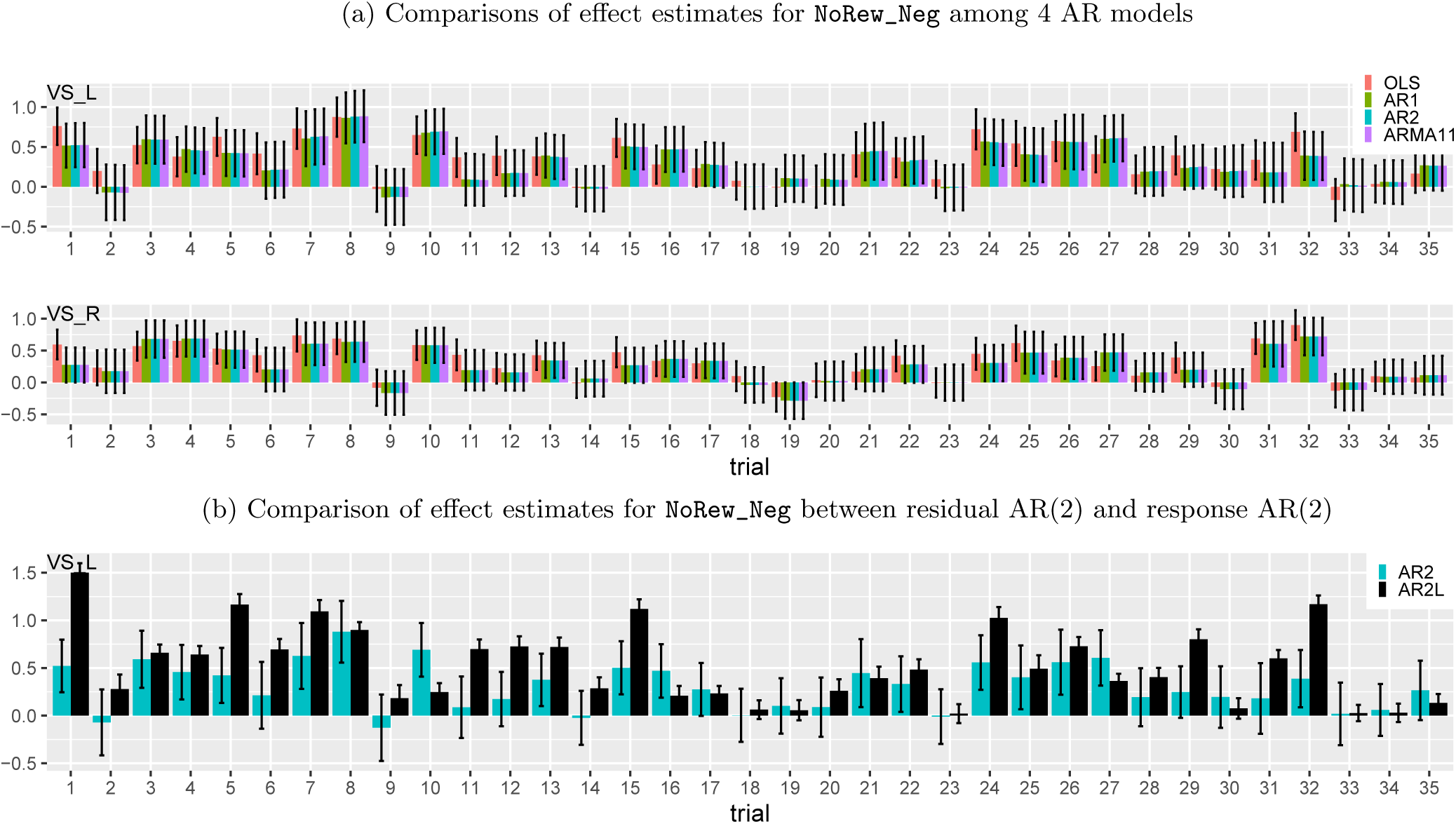
Trial-level effects under the task `NoRew_Neg` from one subject. (a) Effect estimates are shown at two contralateral regions, left (upper row) and right (lower row) ventral striatum. Black segments indicate one standard error, and the colors code the four different AR models (OLS, AR(1), AR(2) and ARMA(1,1)) for the residuals in the GLS model (1). Only 35 trials (out of 48) were successfully completed by the subject. Despite substantial amount of cross-trial variability, some consistent extent of synchronization was revealed across all the four models and all the five contralateral region pairs (only one pair shown here). (b) Effect estimates (AR2L, black) at left ventral striatum were obtained with AR effects modeled as second-order lagged effects of the EPI time series in the model (21) as implemented in Westfall et al. (2017). The same AR(2) results from (a) are shown (AR2, iris blue) as a comparison. The impact of incorporating lagged effects in the model was quite evident with both effect estimates and their precision substantially higher at some trials.

**Figure 14:**
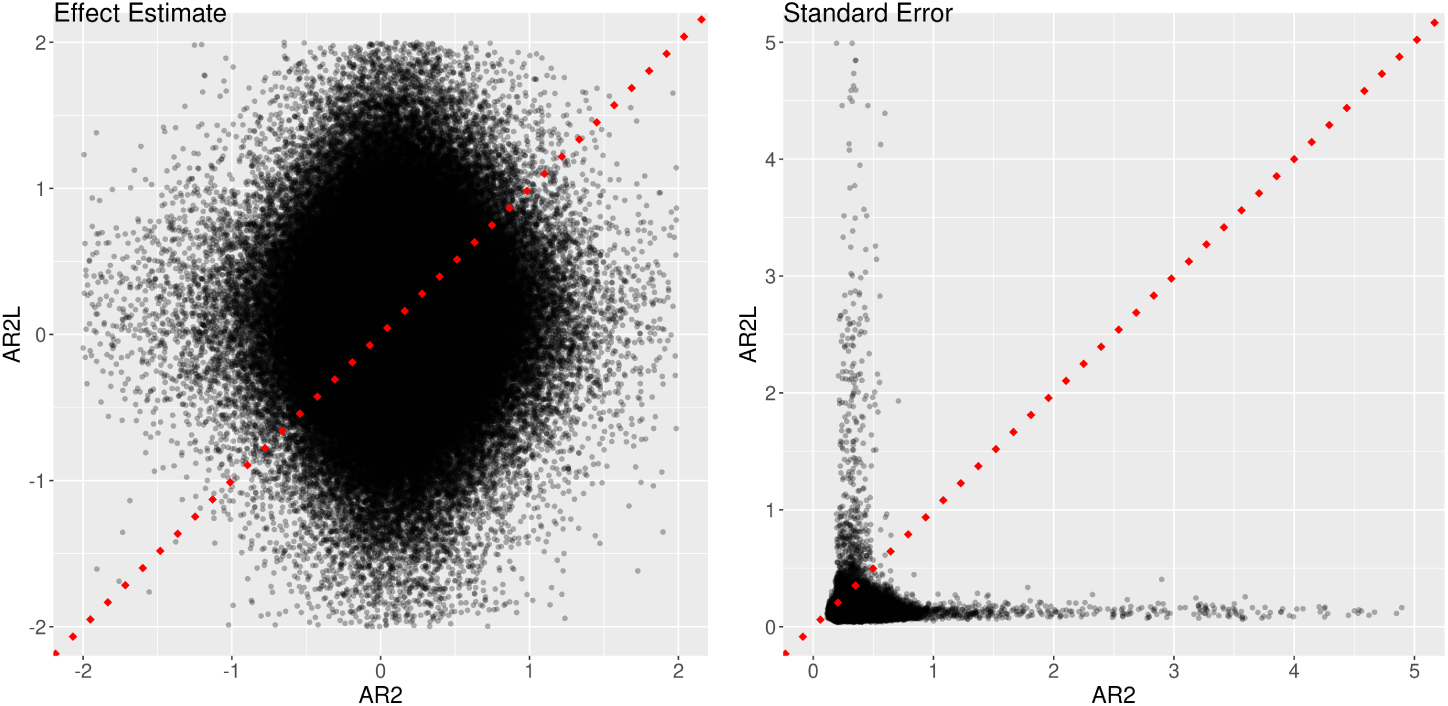
Comparisons of two approaches in AR handling. Two models were adopted to fit the data at the 11 ROIs, one (*x*-axis: AR2) with the GLS model (1) plus an AR(2) structure and the other (*y*-axis: AR2L) with the model (21) that mimicked the approach by Westfall et al. (2017). The effect estimates (left) and their standard errors (right) are shown for the total 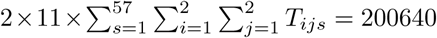 trial-level effects among the two cues and four tasks. The substantial amount of deviation of the effect estimates from the diagonal line (dotted red) indicates the dramatic differences between the two models. The precision underestimation of the model with lagged effects (AR2L) can be assessed by the proportion of data points (98.3%) below the dotted red line.

Among the three AR models, both the AR(2) and ARMA(1,1) models slightly edged out AR(1) due to the extra accountability from the second AR parameter. While a large amount of autocorrelation was explained through the first-order parameters among the three AR models (first, second and fourth columns, Fig. 13), the second parameter for AR(2) and ARMA(1,1) provided less but still sizeable amount of autocorrelation accountability (third and fifth columns, Fig. 13). In light of the observations that both the AR(2) and ARMA(1,1) results were hardly differentiable (Fig. 13), we opted to adopt the ARMA(1,1) model in the current study.

What if the serial correlation is directly modeled as delayed effects of the EPI time series, as adopted in Westfall et al. (2017)? To explore the impact of such approach, we analyzed the EPI data of the 11 ROIs at the subject level with the following model,

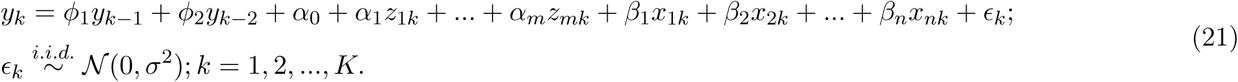

where *k* indexes the time points, *ϕ*_1_ and *ϕ*_2_ are the first- and second-order AR parameters for the lagged effects of the EPI signal *y*_*k*_. As both *y*_*k*−1_ and *y*_*k*−2_ are largely correlated with all the regressors, the impact on effect estimates was substantially evident across subjects, regions, conditions and trials (Figs. 13b,14). In addition to a varying amount of increase on some effect estimates, the uncertainty (standard error) was quite smaller for most effect estimates.

## Footnotes

1 The variabilities across these four levels are usually considered random effects under the conventional statistical framework. In contrast, variations associated with, for example, conditions (e.g. positive, neutral and negative), subject groups (e.g., patients and controls) or quantitative variables (e.g., age, RT), are treated as fixed effects at the population level.

2 Due to the difficulty and varying strategies of handling the discontinuities cross runs and sessions, the analytical pipeline of FMRI analysis can be described in the literature with a two-, three- or even four-level procedure depending on the specific pipeline or software. For example, the analysis for each run may be labeled as the first level, followed by a second level that summarizes the effect estimates across runs (and a third level for across-session summary) through simple averaging or a fixed-effects model; the analysis for generalization at the population level is thus termed as third (or fourth) level. As the data across runs and sessions can be integrated into one model at the subject level through a numerical scheme (e.g., Chen et al., 2012), here we stick to a two-level description for simplification. To avoid the messy terminology in the field, we directly describe the two levels as subject and population instead of their ordinal sequence.

3 The popular term for the subject-level analysis is general linear model (GLM) in neuroimaging. However, a more accurate description of the modeling approach is *time series regression*, especially considering the nature of input data and the complex issue of delicately handling the temporal structure embedded in the residuals through generalized least squares (GLS) (cf., ordinary least squares (OLS) for GLM).

4 The alternative approaches with multiple basis functions share the common assumption of same response magnitude across trials.

5 The program 3dLMEr adopts the same LME framework as its predecessor 3dLME (Chen et al., 2013), but utilizes the *R* package lme4 instead of nlme to accommodate broader modeling capabilities (e.g., handling crossed random-effects structure such as the LME formulation (2)).

6 The i.i.d assumption about cross-trial effects *ηt* in the derivation is likely violated for several reasons including potential serial correlation. However, the overall logic remains applicable.

7 The terms “condition” and “task” are interchangeable in the literature in describing a stimulus type. Here we use “condition” to describe a general category of trials to avoid any potential confusion since the experiment involves both cues and tasks.

8 In general, two factors of *m* and *n* levels, respectively, have an intercept, *m* − 1 and *n* − 1 terms for the individual effects of the two factors, and (*m* − 1)(*n* − 1) interactions. How these total *mn* terms are formulated depends on the specific parameterization method such as dummy and deviation coding.

9 The modeling scripts used in the paper are publicly available at https://github.com/afni-gangc/tlm

10 https://surfer.nmr.mgh.harvard.edu/optseq/

